# Effect of chronic stimulation and stimulus level on temporal processing by cochlear implant listeners

**DOI:** 10.1101/359869

**Authors:** Robert P. Carlyon, François Guérit, Alexander J. Billig, Yu Chuen Tam, Frances Harris, John M. Deeks

## Abstract

A series of experiments investigated potential changes in temporal processing during the months following activation of a cochlear implant (CI) and as a function of stimulus level. Experiment 1 tested patients on the day of implant activation and two and six months later. All stimuli were presented using direct stimulation of a single apical electrode. The dependent variables were rate discrimination ratios (RDRs) for pulse trains with rates centred on 120 pulses per second (pps), obtained using an adaptive procedure, and a measure of the upper limit of temporal pitch, obtained using a pitch-ranking procedure.

All stimuli were presented at their most comfortable level (MCL). RDRs decreased from 1.23 to 1.16 and the upper limit increased from 357 to 485 pps from 0 to 2 months post-activation, with no overall change from 2 to 6 months. Because MCLs and hence the testing level increased across sessions, two further experiments investigated whether the performance changes observed across sessions could be due to level differences. Experiment 2 re-tested a subset of subjects at 9 months post-activation, using current levels similar to those used at 0 months. Although the stimuli sounded softer, some subjects showed lower RDRs and/or higher upper limits at this re-test. Experiment 3 measured RDRs and the upper limit for a separate group of subjects at levels equal to 60%, 80%, and 100% of the dynamic range. RDRs decreased with increasing level. The upper limit increased with increasing level for most subjects, with two notable exceptions. Implications of the results for temporal plasticity are discussed, along with possible influences of the effects of level and of across-session learning.

## Introduction

The processing of electrical stimulation changes in the months following initial activation of a cochlear implant (CI). Probably the most important change from the patient’s perspective is an improvement in speech perception (Blamey et al., 2013), a finding that has been attributed to the patient learning the relationship between the novel form of stimulation and speech segments (Davis et al., 2005). However, other changes, not attributable to learning, also occur. One common observation is that the most comfortable level (MCL), defined as the stimulus level needed to produce a comfortable loudness, increases over time (e.g., Gajadeera et al., 2017). Here we extend the study of plasticity in CI patients to temporal processing, and investigate whether it occurs in adult-deafened human CI patients following activation of their device.

Single-unit recordings from the cat inferior colliculus (IC) have revealed a limitation in the processing of temporal information delivered by a CI. Specifically, neurons in the central nucleus of the IC phase-lock to pulse trains only up to a certain “upper limit” (Snyder et al., 1995; Vollmer et al., 2005; Middlebrooks, 2008; Middlebrooks and Snyder, 2010; Hancock et al., 2013; Vollmer et al., 2017). Importantly, this limit depends on the history of stimulation. For example, juvenile-deafened cats show a higher limit when they have grown up listening through a CI than when they have grown up deaf (Hancock et al., 2013; Vollmer et al., 2017). Evidence for plasticity in adulthood comes from the finding that chronic stimulation as an adult can restore the upper limit of phase locking in cats that have been deaf since birth (Vollmer et al., 2005). There is also some evidence that the upper limit decreases following deafening as an adult. Middlebrooks (2013) found that, in cats deafened as young adults, the upper limit was higher in those that had been deaf a few hours than those that had been deaf for approximately six months. However Hancock *et al* (2013) found no difference in the upper limit between a group of cats that had been deaf for an average of 0.4 months and another group that had been deaf for approximately six months. The different results of the two studies might be due to the different durations of deafness in the shorter-term deafened groups or to other differences in methods and analysis.

Human CI users also show limits in temporal processing. Stimulation of a single or multiple channels with a pulse train causes pitch to increase with increases in pulse rate up to, but not beyond, a certain value. This perceptual upper limit differs across listeners and electrodes, but is typically in the range 200-700 pps / channel (Shannon, 1983; Townshend et al., 1987; Kong and Carlyon, 2009; Kong et al., 2009; Carlyon et al., 2010). It correlates negatively with the duration of deafness, consistent with an effect of auditory deprivation (Cosentino et al., 2016). It also correlates with performance on other tasks that require temporally accurate encoding of high-rate stimuli, including gap detection, and, for bilaterally implanted listeners, the detection of inter-aural timing differences at high pulse rates (Ihlefeld et al., 2015; Cosentino et al., 2016). These correlations cannot be attributed to simple between-subject cognitive differences, such as the ability and/or willingness to concentrate on psychophysical tasks. For example, the upper rate correlates significantly more strongly with gap detection thresholds than with rate discrimination at low rates of about 100 pps (Cosentino et al., 2016), even though the cognitive demands of the three tasks are similar. Furthermore, Ihlefeld *et al* (2015) observed a significant correlation between rate- and ITD-discrimination at high rates across electrodes, using a method that removed across-subject differences. Taken together, the results described above suggest that there may be a sensory limitation that is common to fine temporal processing at high rates, and that is susceptible to auditory deprivation. However, the pattern of results can differ across studies. For example, although Cosentino *et al* did not find a significant correlation between rate discrimination thresholds with a 100-pps standard rate and duration of deafness, such a correlation was obtained by Moore and Carlyon’s (2005) re-analysis of Pfingst’s (1994) data.

The facts that implanted humans and cats appear to show qualitatively similar limitations in temporal processing, and that these limitations are susceptible to experience in the cat, suggests that the perceptual upper limit observed in human CI patients might increase in the weeks and months following activation of their device. Conversely, it is possible that exposure to conventional CI processing strategies, that remove fine timing information, could *reduce* the upper limit. We test whether temporal processing improves or deteriorates in the months following implantation in a study that, in contrast to previous correlational analyses, adopts a longitudinal approach. Specifically, we use two tasks to measure temporal processing at low and at high rates, and do so on the day that the implant is first activated and at two, six, and nine months later. Such results are unaffected by “incidental” differences between subjects and electrodes, such as in the electrode position, local neural anatomy, and cognitive differences – all of which could affect the results of analyses based on correlations. The results revealed a moderate but significant improvement in both tasks during the first two months post switch-on, with an effect size that was greater at high than at low rates. Changes after six months were more variable across subjects with no significant effect overall. Testing at each session was performed at the most comfortable level (MCL), which also increased across sessions. We therefore performed additional control experiments and analyses to determine whether the temporal processing changes could be attributed to these level differences. These include a comparison of performance between the day of activation and nine months later, obtained at similar physical levels (experiment 2), and measurement of the effects of level *per se* with a (different) set of experienced subjects (experiment 3). The results showed that performance generally improved with level, but that some subjects performed better when tested at nine months than on the day of activation, despite the stimuli sounding softer (and having the same physical level).

## 1. EXPERIMENT 1: CHANGES IN TEMPORAL PROCESSING AND MCL BETWEEN 0 AND 6 MONTHS POST-ACTIVATION

### A. Subjects and testing schedule

Nine post-lingually deafened adult users of the Cochlear CI522 implant took part. Their details are shown in Table 1. They were implanted at Addenbrookes hospital, Cambridge, UK. Prior to the activation of the implant they were contacted by a member of the clinical team and invited to participate in the study. The study started with the standard clinical session in which the audiologist measured threshold and comfort levels for each electrode, and identified any electrodes that needed to be de-activated. The processor was then programmed with a clinical map, and the subject listened to live speech for approximately ten minutes. At that point the processor was removed, and the patient undertook an approximately 2-hour session of psychophysical tests during which they made pitch and loudness judgements for pulse trains applied to a single electrode. After a break the patient then undertook a second session with the CI audiologist during which they practiced listening to speech through their processor. The psychophysical session was then repeated after a period of about two months, during which time the patient listened through their device in everyday life but did not perform any psychophysical tests. An additional psychophysical session took place, for the seven subjects still available, approximately six months after implant activation.

**Table 1.** Details of the subjects who took part in experiments 1 and 2. As noted in the text, all subjects were users of the CI522 CI manufactured by the Cochlear company. The duration of deafness is often known only approximately and is rounded to the nearest year. a) For subject C108 a progressive loss was reported 46 years prior to testing, followed by a sudden loss 11 years prior to testing

| Subject | Age at Switch on<br>(years) | Duration<br>of<br>Deafness<br>(years) | Aetiology |
| --- | --- | --- | --- |
| C101 | 70.7 | 16 | Unknown (noise exposure?) |
| C102 | 75.7 | 20 | Otosclerosis, gradual onset |
| C103 | 64.4 | 9 | Unknown, progressive |
| C104 | 54.1 | 54 | Maternal Rubella (congenital) |
| C105 | 74.6 | 48 | Unknown progressive |
| C106 | 58.6 | 58 | Alport's syndrome |
| C107 | 54.4 | 54 | Sudden onset |
| C108 | 73.5 | 46<sup>a</sup> | Unknown progressive |
| C109 | 70.2 | 16 | Unknown progressive |

### B. Stimuli and testing procedure

All stimuli were presented using the NIC3 software and research hardware provided by Cochlear Ltd. This allowed us to bypass the clinical processor and to present trains of symmetric cathodic-first biphasic pulses to electrode 16. All stimulation was in monopolar (MP1+2) mode. The phase duration was 43 μs and the inter-phase gap was 8 μs. The total duration of each pulse train was 500 ms. All stimuli were checked using a test implant and digital storage oscilloscope. Impedances for all electrodes were measured at the start and end of each session, and additionally mid-way through the first session. This was done to ensure that stimulation levels remained within the compliance limits of the device, and as a standard safety check for any increases in impedance that might indicate malfunction. No such increases were observed. Some decreases in impedance were observed during the first (and to a much lesser extent) second session.

The procedure was designed to obtain reliable measures of temporal processing, both at low and high rates, within each of a series of necessarily time-constrained testing sessions. After the initial impedance measure (and check for compliance limits), each session began with measurement of thresholds (T) and Most Comfortable Levels (MCLs) for nine pulse trains, having pulse rates evenly spaced on a logarithmic scale between 90 and 981 pps. Thresholds and MCLs were determined for each subject using loudness scaling. Subjects indicated the loudness of pulse trains using a chart in which loudness was marked on a scale from 0 (‘off’) to 10 (“too loud”). Subjects were asked to listen for a short sound and indicate 1 (“Just noticeable”) on the chart at the first instance they heard a sound. For every subsequent sound, subjects indicated the number that best matched the loudness. For each stimulus, presentation began at 0 current units (CUs), and initially increased in steps of 6 CUs. The level at which the response changed from ‘0’ to ‘1’ was defined as the threshold. Stimulus level was increased and levels corresponding to 2 (‘Very Soft’), 3 (‘Soft’), 4 (‘Comfortable but too Soft’), 5 (‘Comfortable but Soft’) 6 (‘Most Comfortable’) and finally 7 (‘Loud but Comfortable’), were recorded. At point 7, the procedure stopped. The MCL was defined as the mid-point of stimulus levels at which the subject indicated 6 (‘Most Comfortable’). The maximum current step of 6 CUs was reduced to 2 CUs when loudness level ‘4’ was reached. The initial rate tested was 90 pps, followed by 295 pps and 981 pps. Intermediate rates were then measured. Finally, 90pps was tested again, with this second set of measures used for the loudness function (the first set being regarded as a practice run). A locally weighted linear regression model (‘lowess’ in Matlab) was then fitted to this function, and the levels of all stimuli were read off for subsequent testing. This was done to smooth out any slight errors in the loudness function that might occur at one or more rates.

Following loudness estimation we measured rate discrimination thresholds for pairs of pulse trains with rates geometrically centred on 120 pps, using an adaptive procedure (Levitt, 1971). In each two-interval forced-choice trial the subject was required to indicate the interval containing the higher pitch, and the response was scored as correct whenever this corresponded to the higher-rate pulse train. The procedure started with pulse rates of 90 and 160 pps. The rate difference was reduced by a factor of 1.25 after every three consecutive correct responses and increased by the same factor after every incorrect response. Stimulus levels were interpolated from the MCL loudness function for each subject; in the rare cases when the lower pulse rate dropped below 90 pps, which was the lowest rate for which we measured MCL, the level was set to be the same as that for the 90-pps pulse train. This was done because MCLs vary only slightly with rate decreases below about 100 pps (McKay and McDermott, 1998). Correct-answer feedback was provided after every trial. Each change from decreasing to increasing rate difference or *vice versa* was defined as a turnpoint. The step size was reduced to a factor of 1.1 after the first 2 turnpoints. The adaptive run ended after six turnpoints and the threshold, defined as the ratio of the higher and lower pulse rates, was calculated from the geometric mean of the last 4 turnpoints. The adaptive procedure was then repeated and the geometric mean of the two rate discrimination ratios (RDRs) was calculated.

Following the adaptive procedure listeners pitch-ranked eight pulse rates, equally spaced on a log scale between 120 and 980 pps, using the optimally efficient midpoint comparison procedure (“MPC”: Long et al., 2005). This procedure consists of a series of 2IFC trials without feedback. The procedure was run 10 times, each with the stimuli introduced in a different random order, and the pitch rank for each stimulus was calculated from the mean of these 10 “sub blocks”. The pitch-rank function was then fit with a “broken stick”, using the Curve Fitting Toolbox from Matlab. The upper limit of pitch was defined as the rate corresponding to the intersection of two straight lines. To fit the broken stick function, the x-axis values were first transformed to be between 1 and 8, the number of rates. The constraints were respectively [1, 3] and [−0.1, 0] for the slopes of the first and second straight lines, [−10, 1] for the constant term of the first line, and [1 8] for the x-value of the intersection between the two lines. Corresponding start values for the fitting procedure were 1, 0, 0 and 4.5. These fitting parameters were selected by inspecting approximately 120 functions obtained in our laboratory from previous experiments, and choosing a set of parameters that yielded upper limits that corresponded well to visual estimates and that were not unduly affected by occasional outliers. An example of the broken-stick fit is shown for one subject in Fig. 1.

**1).**
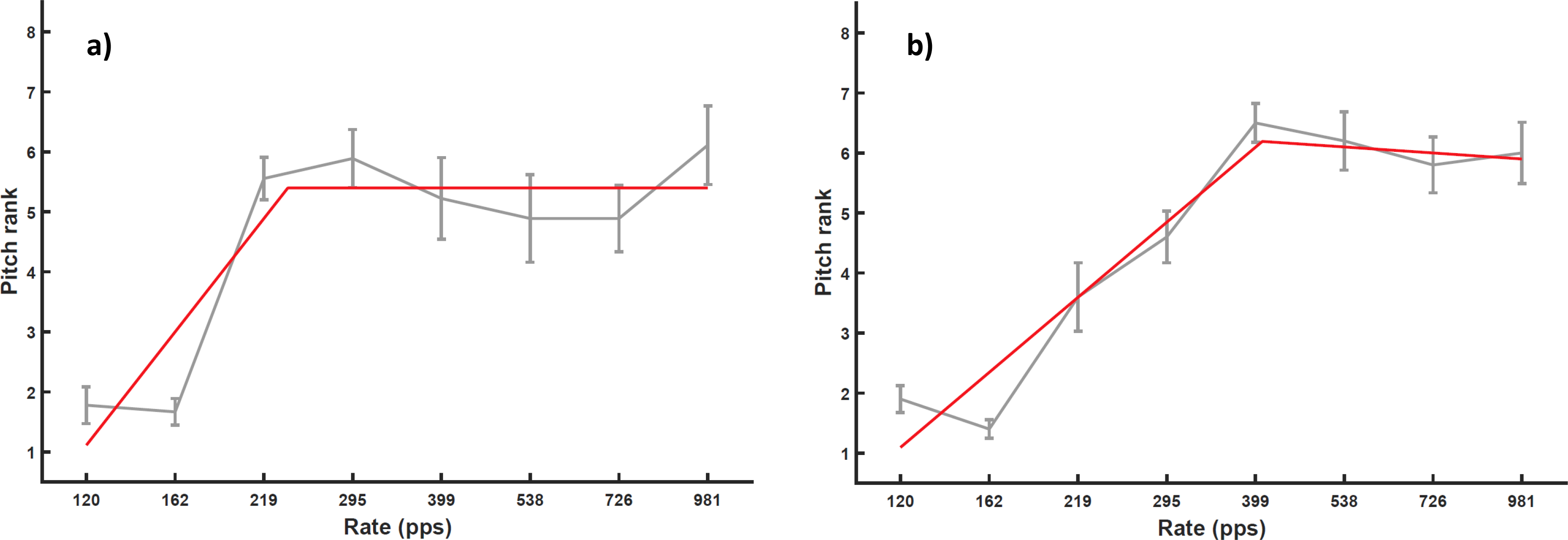
Grey lines show pitch-ranking functions with associated standard errors for one subject, obtained using the midpoint comparison procedure. Red lines show the broken-stick fits to the function. The upper limit is defined as the rate where the two parts of the stick meet. The two plots are for separate sessions.

The low-rate discrimination and pitch-ranking procedures were then repeated. This was done so as to obtain a measure of any initial procedural learning effects, such as might arise, for example, from increased familiarization with the trial structure or from understanding what is meant by the term “pitch”. We assumed that any such effects would be greater within a session than between sessions (during which time the subject listened through their implant but did not perform the psychophysical tests). In contrast, chronic stimulation effects should be larger between than within sessions. Hence greater improvement between than within sessions would be more consistent with an effect of chronic stimulation than of procedural learning. The possible influence of other types of learning will be considered further in the Discussion.

### C. Results

#### Sessions 1 vs 2 (Day of Activation to 2 months)

##### Temporal Processing Measures

Results from the low-rate discrimination measures are shown, for sessions 1 and 2, in two left-hand panels of Fig. 2. Data for individual subjects are shown by the fainter coloured lines; mean data are shown by the solid bold black line. Subject C105 found the task very difficult for all runs and sessions, and, for the first pair of runs in the first session, failed to converge on a threshold. That data point was replaced by the value obtained for the second pair of runs in that session. Even when the procedure converged on a threshold, the large difference between the standard and signal rates means that, for this subject, residual loudness cues may have played a role.

**2).**
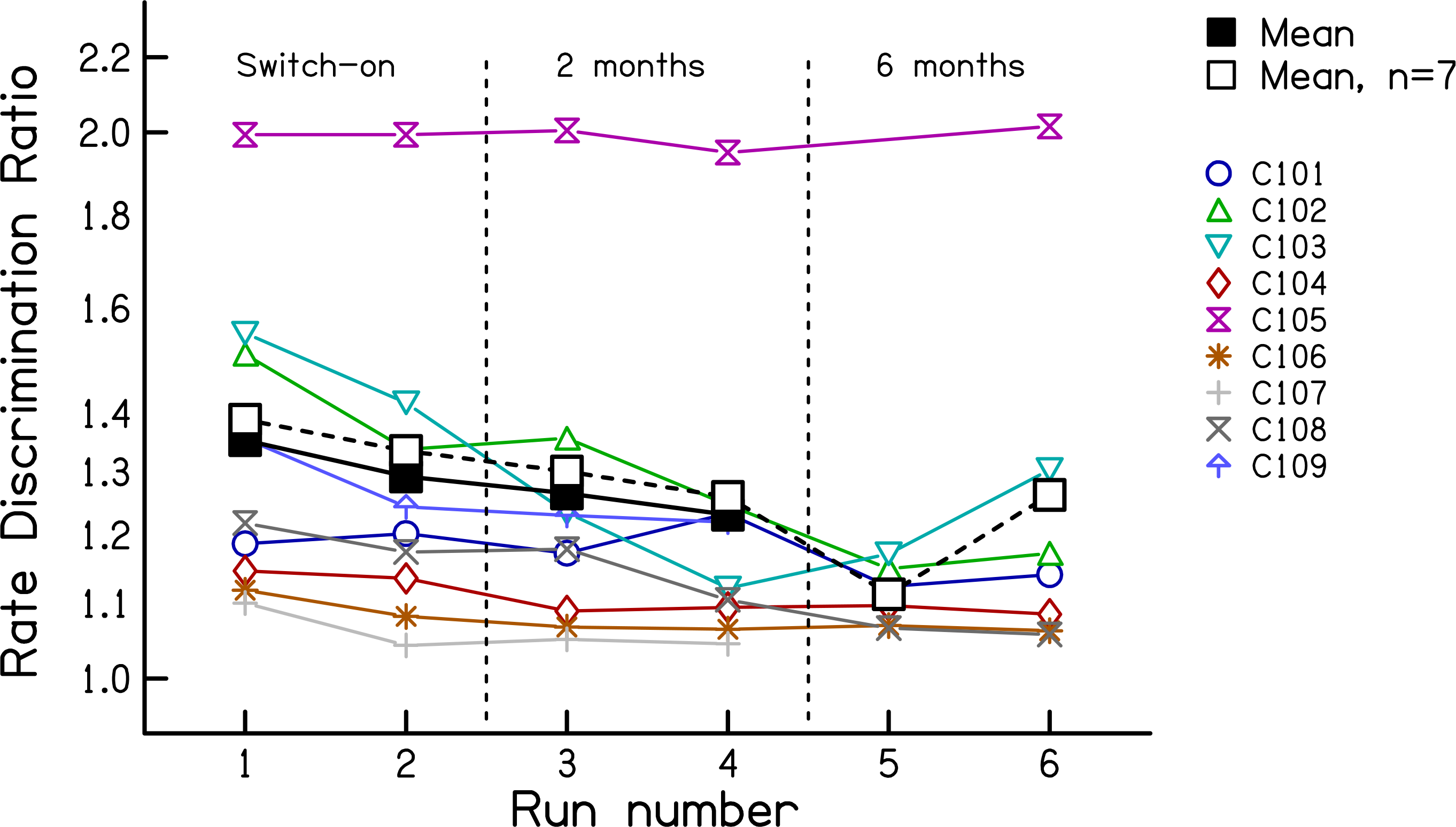
Rate discrimination ratios from experiment 1, measured at 0, 2, and 6 months re the date of implant activation. Coloured symbols show data for individual listeners. The solid black line shows the geometric mean of all subjects. The dashed black line shows the geometric mean of the seven subjects tested at all time points.

The results show a small improvement both within and between sessions. The data were log-transformed and entered into a two-way RMANOVA, with the first vs second pair of runs from each session entered as one factor and session number (activation day vs 2 months) as the other. This revealed main effects of session (F(1,8)=6.11, p=0.039) and within-session run (F(1,8)=6.51, p=0.034); across sessions the RDR decreased from 1.30 to 1.23, corresponding to a factor of 1.06 with 95% confidence limits of 1.005 to 1.120. These effects were also significant when subject C105 was excluded from the analysis (Session: F(1,7)=6.24, p=0.041.Run: F(1,7)=6.12, p=0.043), in which case the RDR decreased from 1.23 to 1.16 across sessions. Note that, for all ANOVAs reported here, we use the Huynh-Feldt sphericity correction and report the uncorrected degrees of freedom.

The upper-limit data are shown in Fig. 3 using the same format as in Fig.2. Note that, opposite to the low-rate RDRs, high values correspond to better performance. A two-way RMANOVA on the log-transformed data revealed a highly significant improvement across sessions (F(1,8)= 33.79; p<0.001) but no effect of within-session run (F(1,8)=0.28, p=0.62). The mean upper limit was 357 pps in session 1 and 485 pps in session 2, corresponding to an increase of 36%, with 95% confidence limits of 20-53%. It is also worth noting that the improvement was significantly larger between the end of session 1 and the start of session 2 than between the two runs of session 1 (t(7)=2.75, p=0.03). This provides some further evidence against the hypothesis that the results were dominated by procedural learning effects.

**3).**
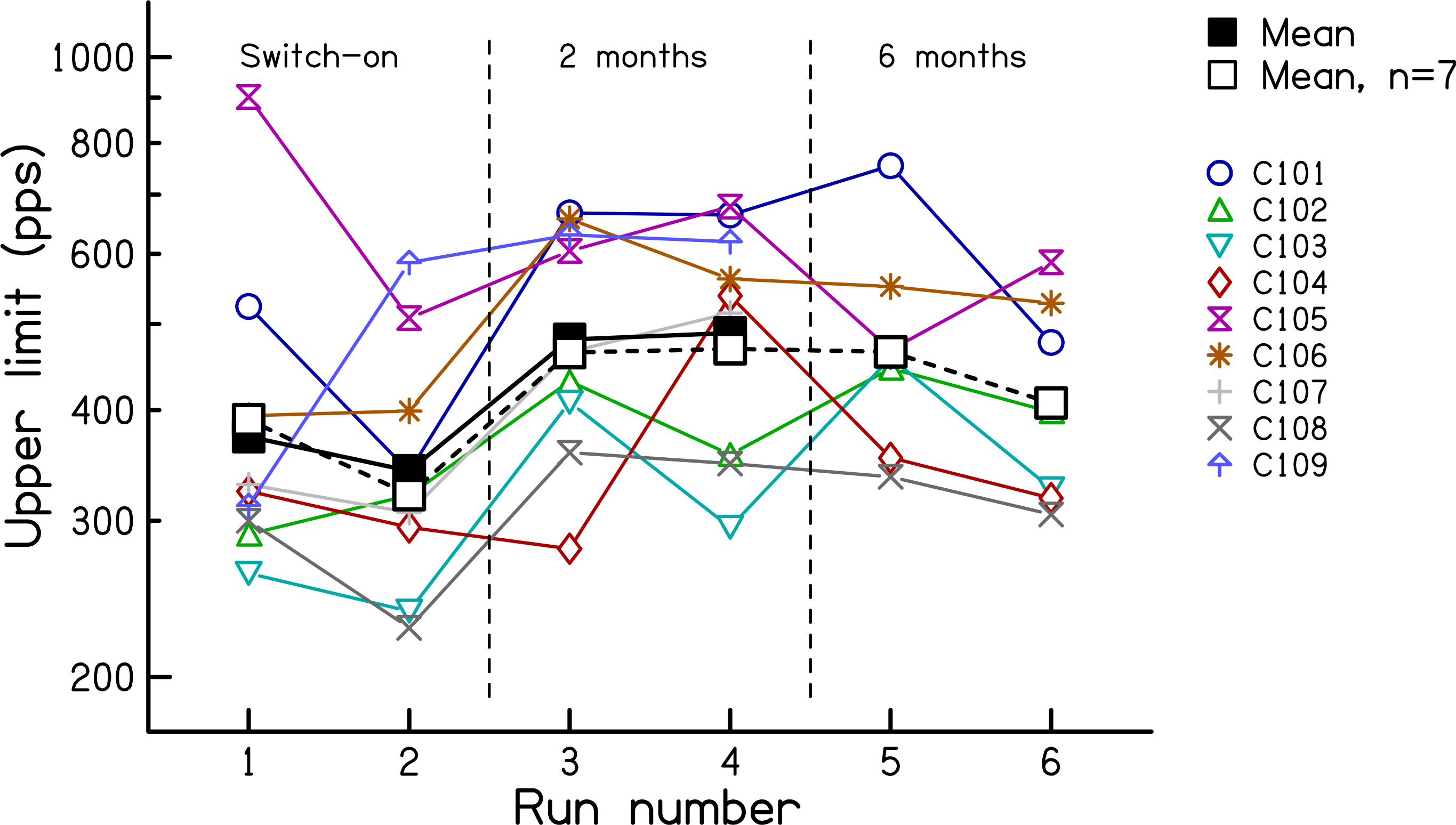
Upper limit measures from experiment 1, measured at 0, 2, and 6 months re the date of implant activation. Coloured symbols show data for individual listeners. The solid black line shows the geometric mean of all subjects. The dashed black line shows the geometric mean of the seven subjects tested at all time points.

To compare the between-session improvements for the two tasks, we calculated the effect size for each subject, again using the log-transformed data. This was initially done separately for each task. For each measure we first calculated the difference between sessions for each subject, and then divided each subject’s difference by the standard deviation of the differences across subjects. This led to a normalized difference for each subject and measure. The sign of the difference was inverted for one measure (because high scores reflect worse performance for the RDR but better performance for the upper limit) and the two measures were then compared using a paired-sample t-test. This revealed that the effect size was significantly larger for the upper limit than for the low-rate RDR (t(7)=2.43, p=0.04).

##### Level differences

As noted above, the upper limit showed a highly significant improvement between sessions, which was significantly larger than that occurring within the first session. Furthermore, the effect size was larger than for the low-rate RDR. Both of these findings are consistent with the idea that chronic stimulation can restore fine temporal processing in a way that is at least partly specific to high-rate stimuli. However, it should be noted that the stimulus level was higher in session 2 than in session 1, due to the increase in MCL between the two sessions (RMANOVA: session F(1,8)=14.44, p=0.005; rate F(8,64)=27.79, p<0.001; session X rate F(8,64)=0.49, p=0.55). That difference is shown by the coloured lines in Fig. 4 for each subject, with the mean data shown by the thicker black line. The difference, averaged across subjects and rates, amounted to 1.7 dB. In experiment 3 we show that, for a different group of listeners, a level difference of approximately this size could produce an increase in the upper limit of about 29-44%, and that the size of this difference did not differ significantly from that observed here between sessions 1 and 2.

**4).**
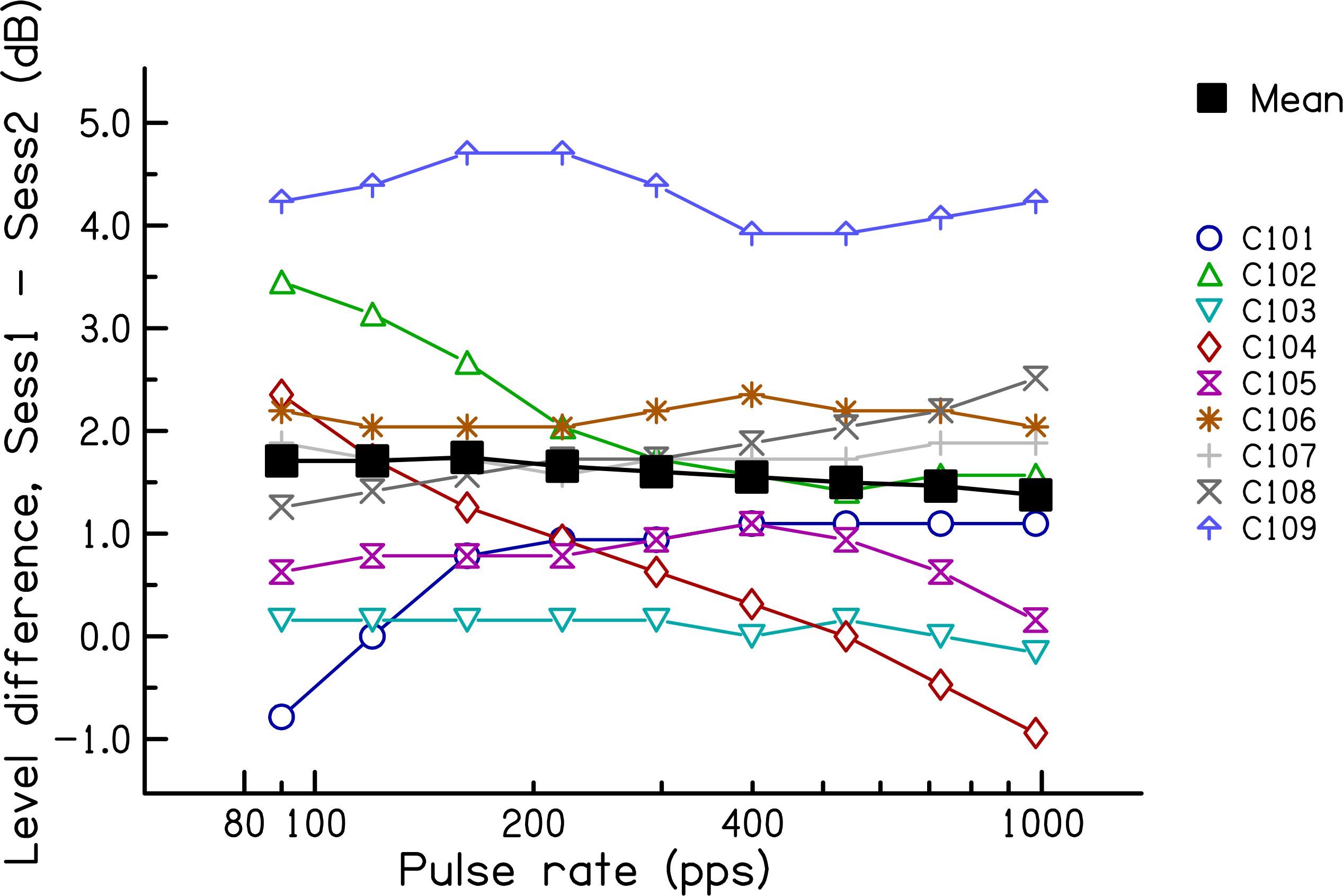
The difference in stimulus level between the 2- and 0-month sessions of experiment 1, plotted as a function of pulse rate. Coloured symbols show data for individual listeners. The solid black line shows the geometric mean of all subjects.

Further evidence for a level-based explanation would occur if the level and upper-rate differences between sessions correlated positively across subjects. However, this correlation was small, negative, and not significant (r(7)=−0.30, p=0.44). It nevertheless remains possible that the between-session improvement was mediated by the change in level. Because the level change was similar at all rates, but the effect size was larger for the upper limit than for the low-rate RDRs, this explanation would require that the upper limit was more susceptible than the low-rate RDR to differences in level. This and other issues relating to the possible effects of level are addressed in experiments 2 and 3.

#### Sessions 2 vs 3 (2 vs 6 months after activation)

##### Temporal Processing Measures

Low-rate RDRs for the seven subjects re-tested at 6 months are shown in the right-hand panel of Fig. 2. The mean data for those seven subjects are shown by the bold dashed line and open squares. There was no significant difference either between the RDRs obtained at 2 vs 6 months (F(1,6)=1.36,p=0.29), or between the two runs in each session (F(1,6)=0.78 p=0.41).

The upper limit data are shown in the right-hand panel of Fig. 3, using the same format as in Fig. 2. There was no significant difference either between the upper limits obtained at these two time points (F(1,6)=2.57,p=0.16), or of within-session run (F(1,6)=0.57 p=0.48).

##### Level differences

Fig. 5 shows the level difference in sessions 2 and 3 relative to that in session 1, averaged across all rates. Data for each subject are shown by the faint coloured lines; mean data are shown by the solid black line. Levels were generally higher in session 3 (6 months post switch-on) than in session 2 (2 months), but this difference just failed to reach significance (t(6) = 2.37, p=0.055). The level difference varied across subjects, but was substantial for some subjects. The largest level increases occurred for subjects C105 (2.0 dB) and C106 (2.3 dB). Note that neither of these two subjects showed a substantial increase in the upper limit between these two sessions (Fig. 3).

**5).**
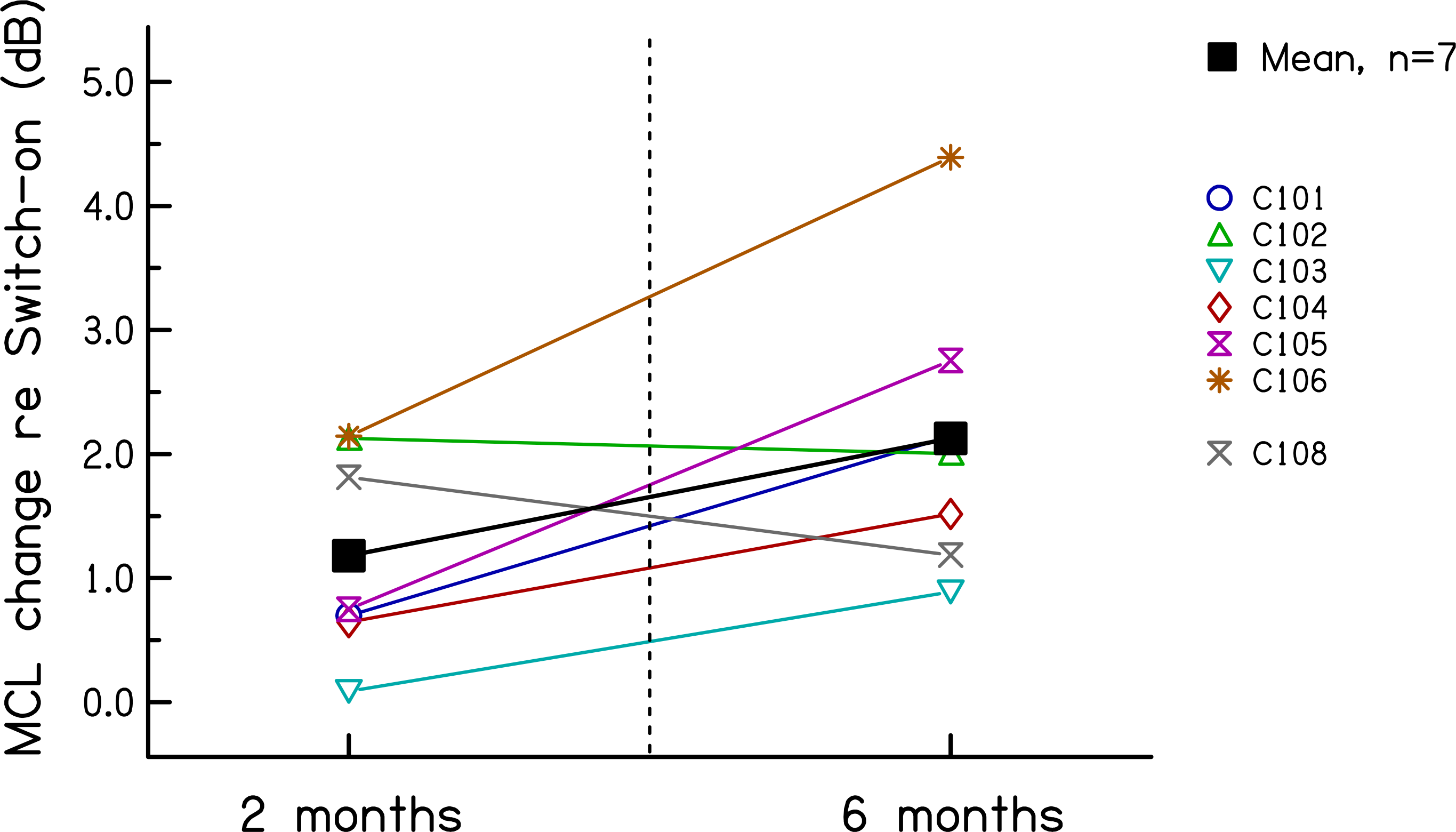
Difference in MCL, averaged across rates, at 2 and 6 months relative to that measured at 0 months in experiment 1.

### D. Summary

Both the upper limit and the low-rate RDR improved from 0 to 2 months post-activation, but not between 2 and 6 months. There was no evidence for a decrease in the upper limit, as predicted if the brain “learns” to ignore fine temporal cues. Instead the upper limit increased by an average of 36% during the first two months. The moderate size of this improvement imposes an upper constraint on the amount of plasticity in temporal processing that occurs during the first two months after implant activation. Some or all of this improvement might arise from the increase in MCL and hence testing level between 0 and 2 months. Strands of evidence against this latter interpretation are that a) the effect size was larger for the upper limit than for the low-rate RDR, whereas the MCL changes were similar across rates, b) there was no across-subject correlation between level-and upper-limit differences between 0 and 2 months, and c) some subjects showed an increase in MCL but no increase in upper limit between 2 and 6 months. We re-visit this evidence in the Discussion, and the possible influence of level differences on the results obtained in the first two sessions is examined further in experiments 2 and 3.

## 2. EXPERIMENT 2: TEMPORAL PROCESSING AT 0 VS 9 MONTHS, EQUAL LEVEL

As noted above, both of the measures of temporal processing and the MCL increased between the day of activation and two months later. To shed further light on whether temporal processing *per se* improves after chronic stimulation, the six remaining available subjects from experiment 1 were re-tested on the upper limit and low-rate discrimination tasks using similar physical levels to those employed on the day of activation. This was done approximately nine months after activation. Note that, due to the change in MCL, these same levels will have sounded softer than on the day of activation. This additionally provides a unique opportunity to, at least partially, distinguish between the effects of level and loudness; for non-longitudinal studies, varying current level for the same stimulus is necessarily accompanied by a change in loudness.

### A. Method

The methods were similar to those employed in experiment 1, with the exception of the setting of the stimulus levels. This was done in order to use levels as similar as possible to those employed on the day of activation, with the constraint that loudness should be similar across rates. We first obtained loudness growth functions for each pulse rate using the same loudness scaling chart as in experiment 1, and then read off (using linear interpolation when necessary) the loudness scaling values corresponding to the stimulus levels used on the day of activation. We then calculated the average of these loudness values across pulse rate. Next, we used the loudness growth function for each pulse rate to calculate the stimulus level needed to elicit this average loudness value. Finally, the current levels were smoothed using a locally weighted linear regression function as described for experiment 1.

### B. Results

The level differences between the 9- and 0- month data are shown for each subject, as a function of rate, in Fig. 6. It can be seen that the differences are, as planned, on average very small. Averaged across all subjects and rates, the level was 0.1 dB higher in the later session. This difference was not significant. Some subjects showed rate-dependent level differences. For subject C104, the levels used at the 9-month point were higher than those on the day of activation at low rates, but lower at high rates. The opposite pattern was observed for subject C106 and, to a lesser extent, C101.

**6).**
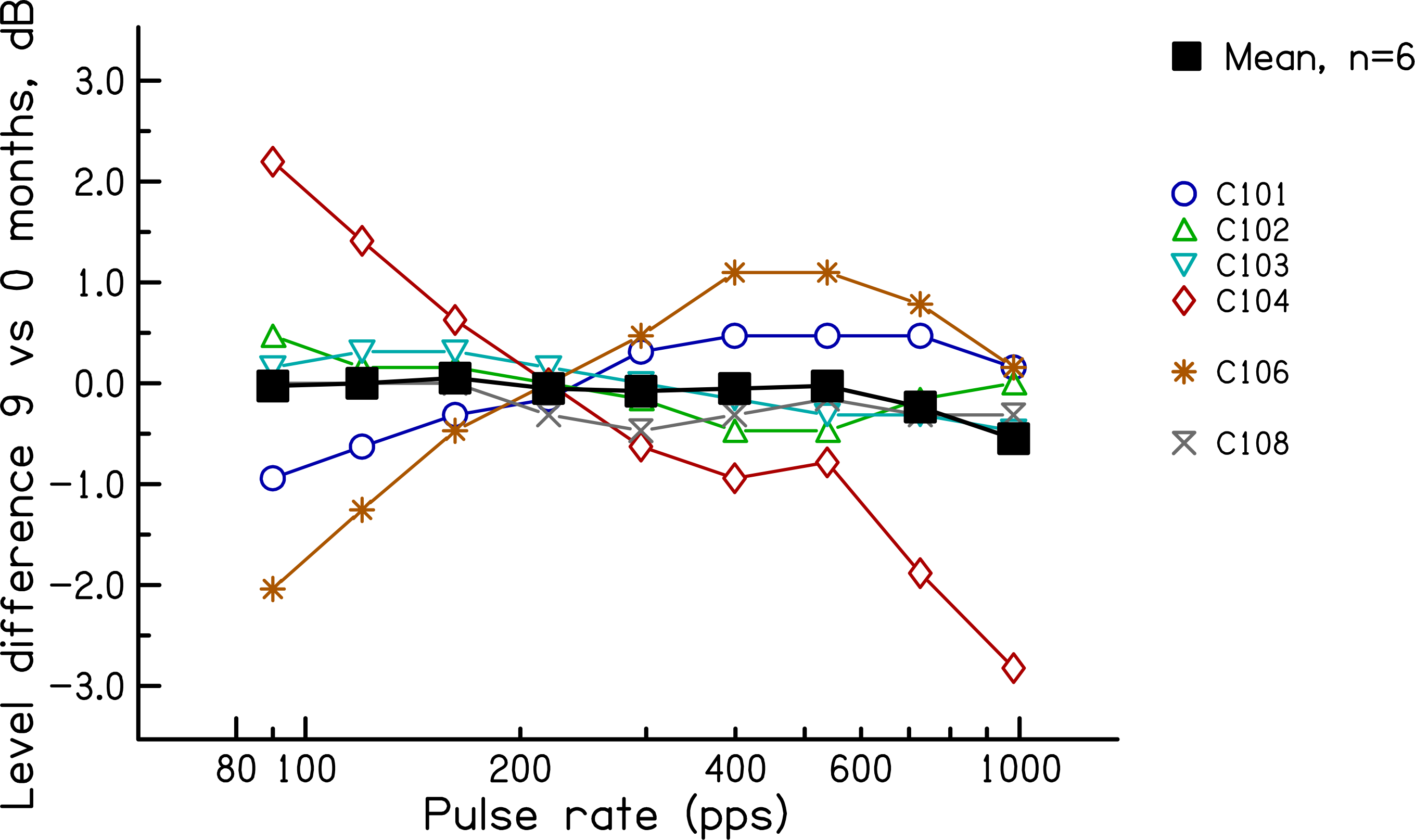
Level difference between the stimuli used at 9 and at 0 months after the day of activation, as a function of pulse rate. Coloured symbols show data for individual listeners. The solid black line shows the geometric mean of all subjects.

The low-rate RDRs are shown for each subject in Fig. 7. The data were averaged across all runs within a session. The average loudness value across pulse rates at nine months is given on the abscissa, next to each subject’s identifier. Recall that on the day of activation this level was 6, corresponding to “most comfortable”. At nine months the values were, for different subjects, roughly 2 (“very soft”, C106), 3 (“soft”, C108) and 4 (“comfortable but too soft”, all other subjects). As a group the RDRs did not differ significantly between the two sessions (t(5)=1.38, p=0.23). However, some subjects seem to show a difference between the two sessions. To assess this we performed, for each subject, independent-sample t-tests on the four runs obtained for each session. This revealed significant differences for all subjects except C101; all of these survived correction for multiple comparisons using the Holm-Bonferroni correction (Holm, 1979). For these subjects, all differences were in the direction of lower RDRs on the later session, with the exception of C106 who was tested at a “very soft” loudness level of 2.2 in that session. The increase in RDR for C106 is evidence that loudness, rather than simply the subject’s report of loudness, had genuinely decreased over nine months and that this impaired performance.

**7).**
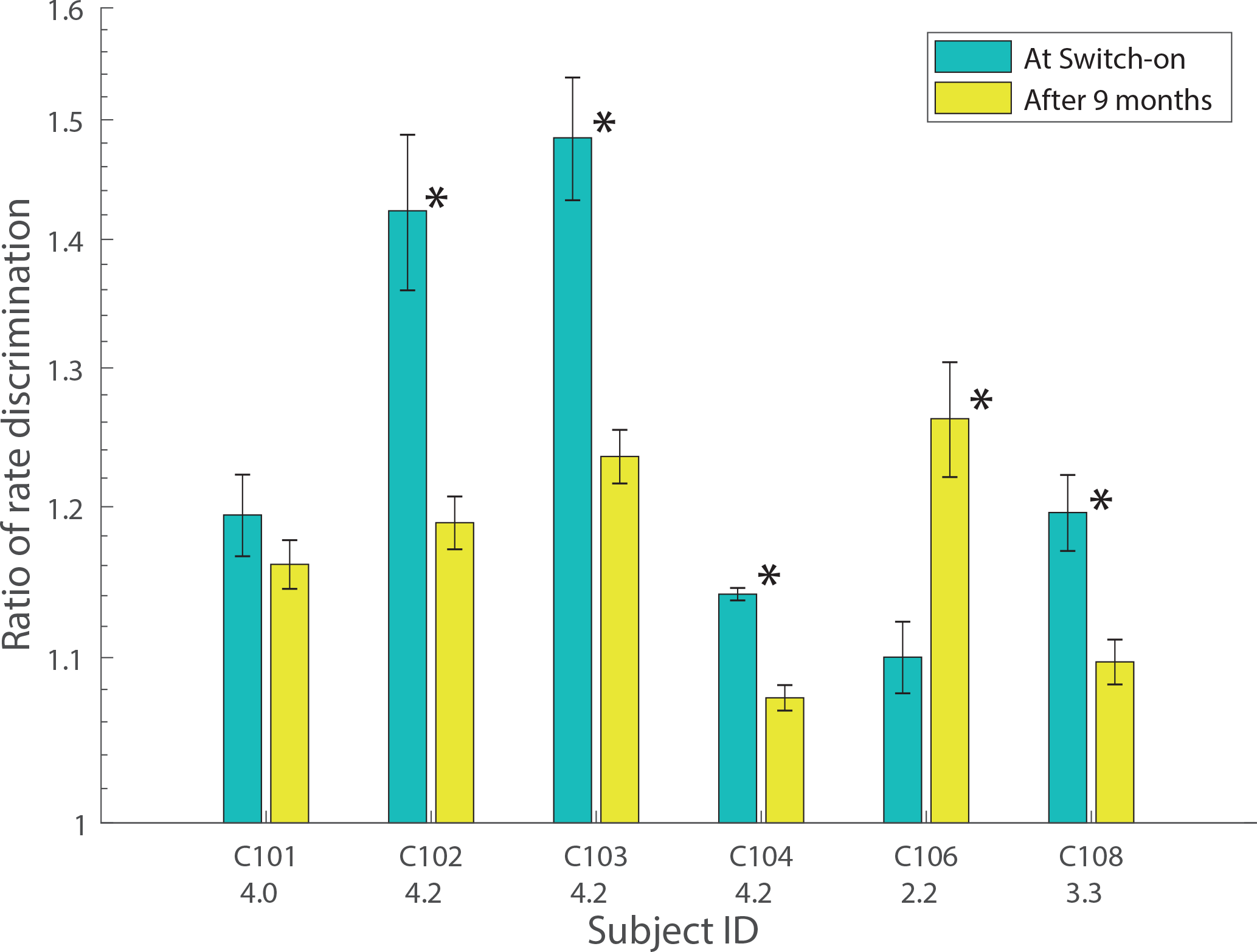
RDRs measured at 0 and 9 months after the day of activation, for six subjects. Significant differences between the two sessions are indicated by asterisks. Error bars show the standard error across runs. The number underneath each subject’s identifier on the abscissa shows the average loudness rating for the 0-month stimuli when judged at 9 months (see text for details).

The upper-limit data are shown in Fig. 8 using the same format as in Fig. 7. These values were obtained by averaging the upper limits obtained from the first and second block of 10 runs in each session. To test whether any of the observed differences were statistically significant, we performed the following analyses. First, from the 20 sub-blocks in each session we obtained 200 samples of 10 sub-blocks, from each of which we derived a broken-stick estimate of the upper limit. The average of these values was taken as the upper limit for each session; it was always within 8% of that obtained by averaging the MPC from the first and last 10 blocks and shown in Fig. 8. The difference between these upper limits (session 4 – session 1) is shown for each subject by the vertical line in the corresponding panel of Fig. 9. We then combined all 40 MPC sub-blocks (20 from each session) and, for each of 200 samples, calculated the difference between the upper limits estimated from two randomly selected sets of 10 sub-blocks. The purpose of this last step was to estimate the distribution of differences that would occur under the null hypothesis that the upper limits observed in sessions 1 and 4 arose from the same underlying distribution. These null distributions are shown by the green bars. As expected they are all centred on a value close to zero.

**8).**
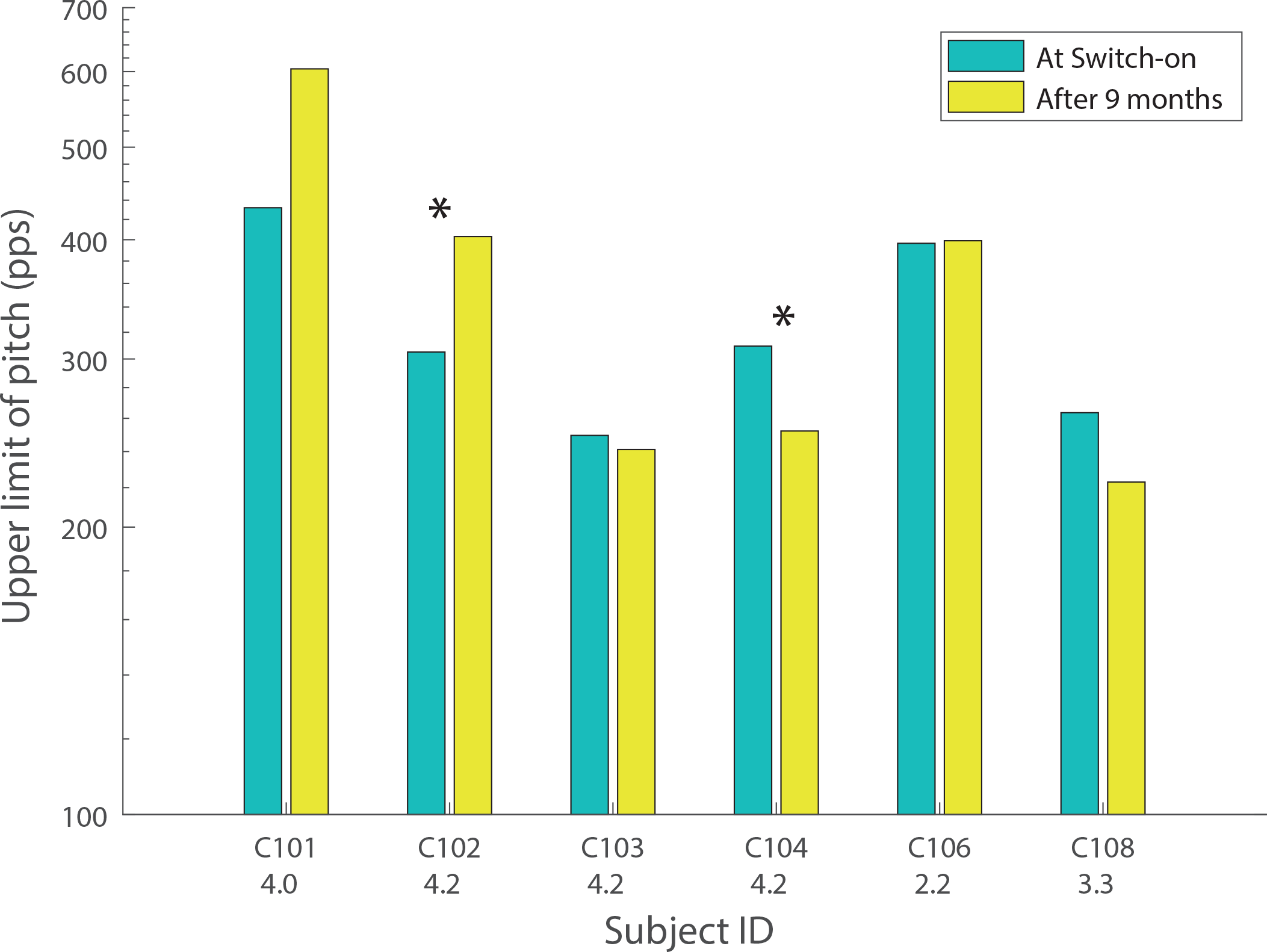
Upper limits measured at 0 and 9 months after the day of activation, for six subjects. Significant differences between the two sessions are indicated by asterisks. The number underneath each subject’s identifier on the abscissa shows the average loudness rating for the 0-month stimuli when judged at 9 months (see text for details).

**9).**
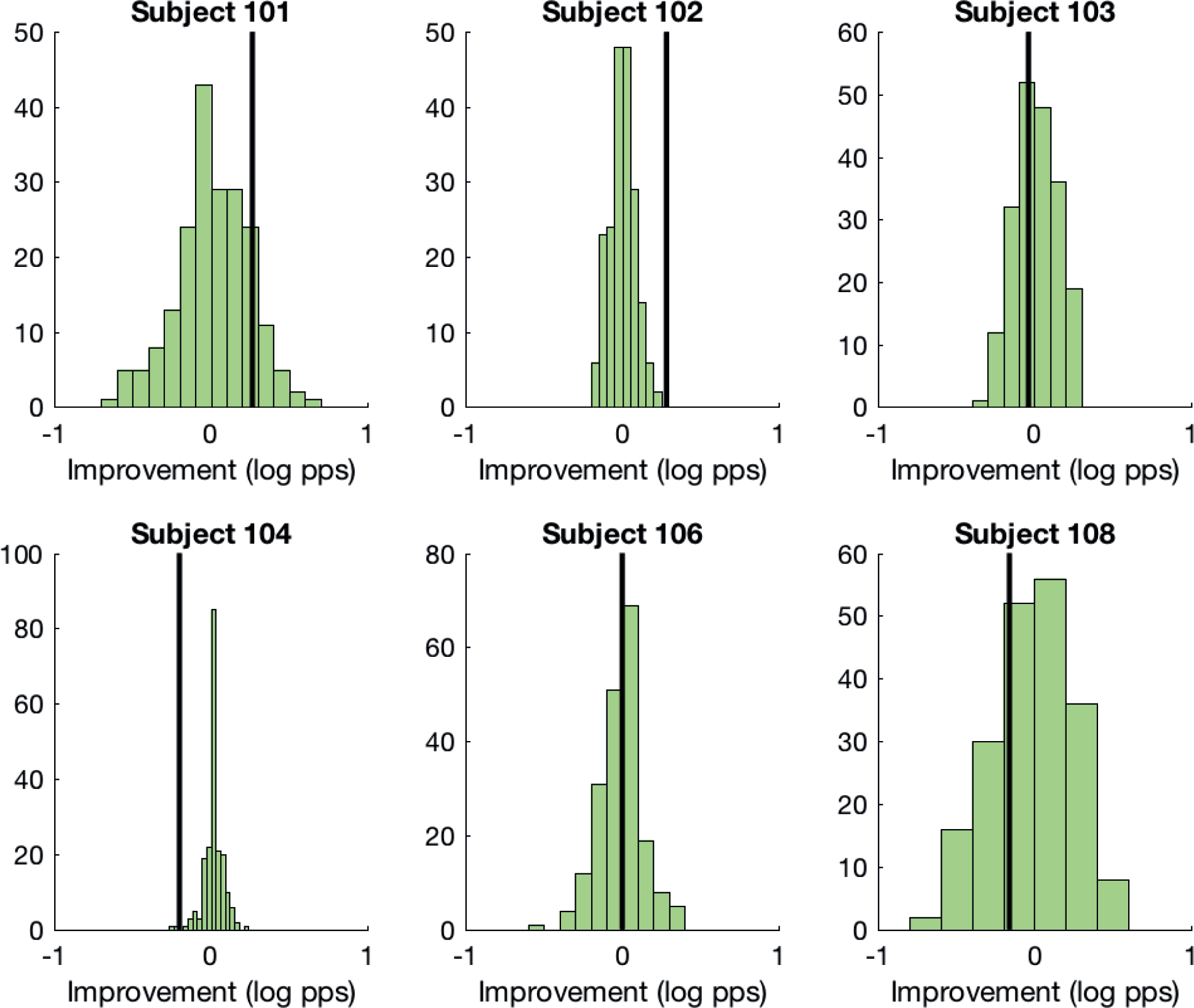
The vertical lines in each plot show the change in the upper limit when measured at 9 months, compared to 0-month measurements obtained at a similar stimulus level. The green histograms show the distribution of differences obtained under the null hypothesis of no difference between the two sessions. (See text for details).

For two of the subjects (C102 and C104), the observed value (vertical line) fell outside of the range of the vast majority of simulated trials. For subject C102 the difference between sessions 4 and 1 was greater than zero, and was also greater than that obtained from all of the 200 simulated trials. For this subject, then, the upper limit increased from session 1 to session 4, despite the softer loudness (and same physical level) in the latter session. This is consistent with a beneficial effect of chronic stimulation. It could conceivably be due to a procedural learning effect, although we note that this subject’s upper limit did not change appreciably during the first two halves of the first session. Subject C104 showed the opposite effect: the difference was less than zero and lower than 197 of the 200 simulated runs. It is possible that this difference arose because, although the average level was similar across rates, the higher rates – which are likely to be most important for the upper limit-had a lower level on session 4 than on session 1 for this subject (Fig. 6). The solid lines in Fig. 10 show the raw MPC functions for sessions 1 (red) and 4 (blue) for subjects C102 and C104; the corresponding broken-stick fits are shown as dashed lines. These MPCs, which were obtained from the average of all 20 sub-blocks in each session, illustrate the improvement for subject 102 and worse performance for subject 104 in session 4 compared to session 1.

**10).**
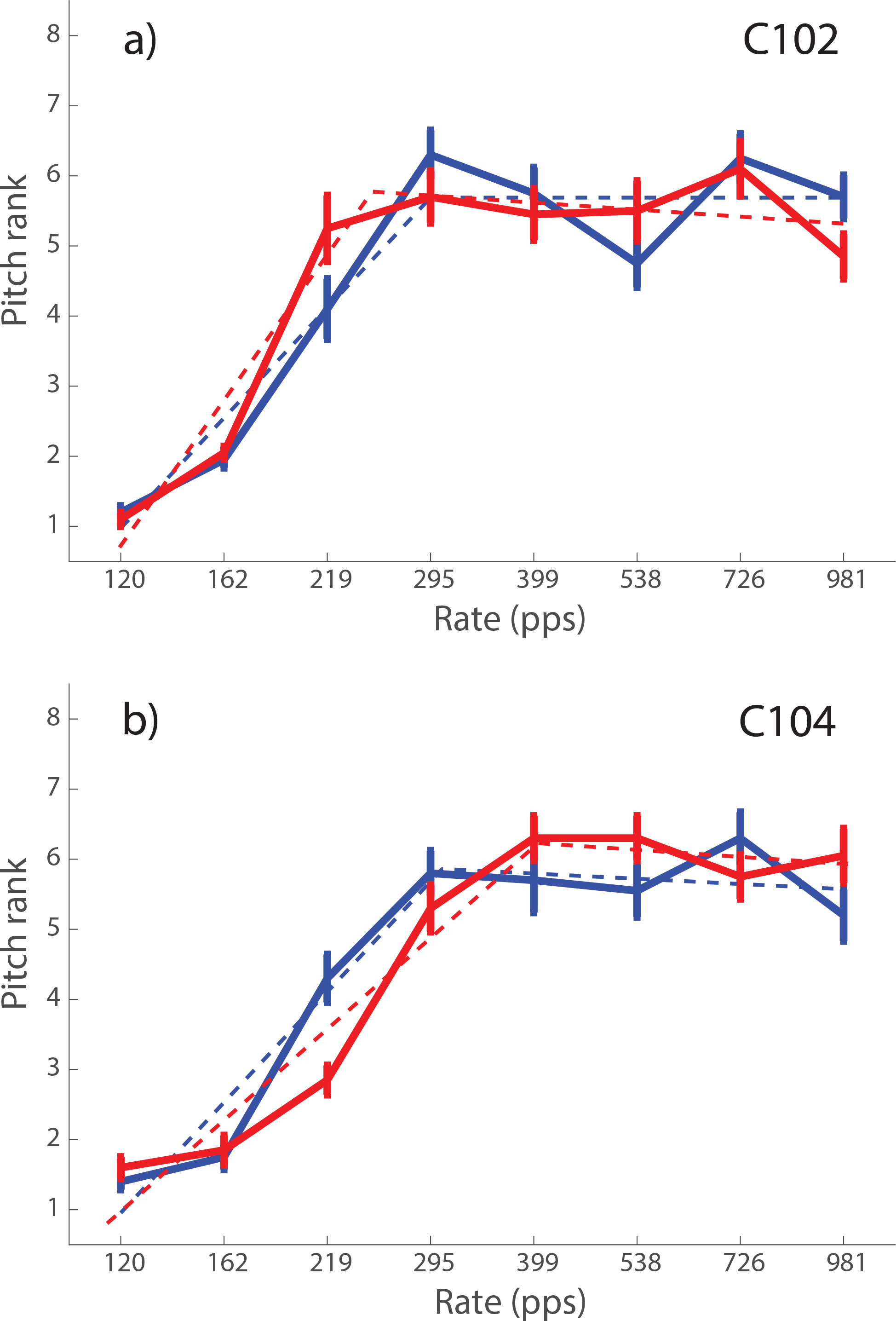
Blue and red solid lines show the pitch rank functions obtained at 0 and 9 months post activation, respectively, for stimuli of similar levels. The dashed lines show broken stick fits. Each panel shows data for one subject.

Finally, our results provide further support for the conclusion reached by Cosentino *et al* (2016), which is that the temporal processing limitations revealed by the upper limit measure seem to have a different basis from those revealed by the low-rate RDR measure. As in the Cosentino *et al* study we observed good test-re-test reliability for each measure in isolation, but no across-subject correlation between the two tasks. To evaluate this, we first measured the across-subject correlation between sessions 1 and 2, which was 0.93 for the RDR and 0.86 for the upper limit, both of which were highly significant (t(7)=8.86, p<0.001 and t(7)=4.46, p<0.005 respectively). We then measured the correlation between the two tasks, averaged across sessions, which should be negative if the across-subject variation in performance on the two tasks was dominated by a common limitation. However, the correlation was positive (r=0.46), mostly due to the fact that subject C105 had a very high RDR and also a high upper limit. When C105’s data were excluded the correlation was slightly negative and not significant (r=-0.248, t(7)=0.68, p=0.52).

## 3. EXPERIMENT 3: TEMPORAL PROCESSING AS A FUNCTION OF LEVEL

To gain further insight into the effects of level on temporal processing, experiment 3 measured the upper limit and low-rate discrimination as a function of level. Because the subjects from experiments 1 and 2 had committed to a limited number of testing sessions, a different set of 11 subjects took part in experiment 3. These subjects were all experienced in the two psychophysical tasks under test, and used a variety of devices. Each subject had typically performed a previous temporal processing task using the electrode selected for this study. Subject details, along with the electrode tested for each subject, are given in Table 2. One MedEL user who was initially tested using electrode 2 was additionally re-tested using electrode 6, leading to a total of twelve data sets.

**Table 2.** Details of the subjects who took part in experiment 3. Note that subject ME5 was re-tested on electrode 6, and, for that electrode, is designated as subject ME5_b

| Subject | Manufacturer | Device | Electrode array | Processing strategy | Electrode tested | Age at testing (years) | Duration of CI use (years) |
| --- | --- | --- | --- | --- | --- | --- | --- |
| C1 | Cochlear | CI24RE | Contour Advance | ACE | 11 | 75.8 | 8.9 |
| AB1 | Advanced Bionics | HiRes90k | Hi Focus 1J | HiRes-S | 8 | 72.2 | 6.9 |
| AB6 | Advanced Bionics | HiRes90k | Hi Focus 1J | HiRes-S | 3 | 68.6 | 2.8 |
| ME5 | Medel | Concerto | FLEX28 | FS4 | 2 | 59.6 | 3.6 |
| AB2 | Advanced Bionics | HiRes90k | Hi Focus 1J | HiRes-S | 8 | 58.1 | 8.4 |
| C8 | Cochlear | CI24RE | Contour Advance | ACE | 11 | 70.4 | 11.0 |
| AB26 | Advanced Bionics | HiRes90k | HiFocus ms | HiRes-S | 3 | 56.1 | 1.9 |
| C3 | Cochlear | CI24RE | Contour Advance | ACE | 11 | 73.5 | 10.3 |
| C9 | Cochlear | CI24RE | Contour Advance | ACE | 11 | 67.1 | 10.3 |
| ME5_b | Medel | Concerto | FLEX28 | FS4 | 6 | 59.7 | 3.7 |
| ME2 | Medel | Concerto | FLEX28 | FS4 | 2 | 50.5 | 3.5 |
| ME15 | Medel | Concerto | FLEX28 | FS4 | 2 | 36.8 | 3.1 |

**Table 3.** Level differences in dB, relative to those used at 100% of the DR, for the 80%- and 60%-DR conditions of experiment 3. Each column shows the level difference for one pulse rate and for different subjects. The subject codes are the same as in Table 2.

| Loudness Balanced Levels, Re: 100% DR (dB) |  |  |  |  |  |  |  |  |
| --- | --- | --- | --- | --- | --- | --- | --- | --- |
|  | 100% - 80% |  |  |  | 100% - 60% |  |  |  |
|  | 90 | 162 | 399 | 981 | 90 | 162 | 399 | 981 |
| C1 | 1.412 | 1.098 | 1.255 | 1.725 | 2.824 | 2.510 | 2.980 | 4.078 |
| AB1 | 0.811 | 0.820 | 0.807 | 1.344 | 1.647 | 1.305 | 1.348 | 2.813 |
| AB6 | 1.153 | 0.905 | 0.946 | 1.513 | 2.302 | 2.034 | 2.197 | 2.917 |
| ME5 | 1.101 | 1.101 | 1.074 | 1.235 | 2.227 | 2.095 | 2.168 | 2.258 |
| AB2 | 1.167 | 1.237 | 1.059 | 0.515 | 2.323 | 2.205 | 1.624 | 1.460 |
| C8 | 1.569 | 1.098 | 1.725 | 2.039 | 3.294 | 2.510 | 2.980 | 3.765 |
| AB26 | 1.053 | 0.955 | 0.483 | 0.698 | 2.012 | 1.925 | 1.467 | 2.570 |
| C3 | 1.569 | 1.098 | 1.098 | 1.412 | 2.980 | 2.667 | 2.510 | 2.824 |
| C9 | 2.196 | 2.196 | 1.882 | 1.726 | 4.235 | 3.294 | 2.824 | 2.980 |
| ME5_b | 0.948 | 0.728 | 0.662 | 0.593 | 1.920 | 1.615 | 1.568 | 1.421 |
| ME2 | 0.915 | 0.720 | 0.828 | 0.848 | 1.818 | 1.623 | 1.743 | 1.913 |
| ME15 | 1.122 | 1.122 | 1.253 | 1.404 | 2.239 | 2.071 | 2.370 | 2.718 |
| Mean | 1.251 | 1.090 | 1.089 | 1.254 | 2.485 | 2.154 | 2.148 | 2.643 |

### A. Method

The same general method was used for all subjects, regardless of the device they were implanted with. In particular, we used a dB scale to adjust current levels and to calculate dynamic ranges for all subjects, even though current level is usually specified in linear units for the MedEL and Advanced Bionics devices. As in experiments 1 and 2, all stimuli consisted of 500-ms trains of symmetric cathodic-first biphasic pulses applied to a single electrode. The same Matlab program was used to control stimulus presentation and record responses for all three devices. The program called low-level routines provided by the respective implant manufacturers as part of the NIC3 (Cochlear), RIB2 (MedEl) and BEDCS (Advanced Bionics) software packages. Stimuli were identical to those used in Experiment 1, except that Advanced Bionics and Medel devices had an inter-phase gap of 0 μs rather than the 8 μs used for the Cochlear device.

We first measured MCLs for pulse rates of 90, 162, 399, 981 pps in ascending order, followed by a re-measurement of the MCL at 90 pps to check for order effects. These measurements were obtained using the same loudness chart and adjustment method as in experiment 1. We then measured detection thresholds at 90 pps using a 2IFC trial structure and an up-down adaptive procedure (Levitt, 1971), so that we could specify stimulus levels in terms of the percentage of dynamic range for the temporal tasks. The stimulus level was initially well above threshold and below MCL. Its level was decreased after every three correct responses and increased after every incorrect response. The change from decreasing to increasing level defined a turnpoint. The step size was 2 dB for the first two turnpoints and 1 dB thereafter. The procedure stopped after eight turnpoints and threshold was estimated from the mean of the last six turnpoints. Two adaptive runs were made, and the average taken of the two.

We then performed loudness balancing, starting with the 90-pps pulse train at its MCL as a reference and the 160-pps pulse train as the variable stimulus. The subject adjusted the level of the variable stimulus so that its loudness was judged equal to that of the reference stimulus, and was encouraged to bracket this equal-loudness level several times before their final adjustment. The 160-pps pulse train, at this adjusted level, then became the reference stimulus and the 90-pps pulse train was loudness-matched to it in the same fashion. The average of the level difference between the 90- and 160-pps pulse trains was then used to set the level of the 160-pps pulse train, which was defined as its new MCL. We also sometimes refer to this level as 100% of the dynamic range (“DR”). This procedure was then repeated so as to match 399 to 162 pps, and 981 to 399 pps. This completed the loudness balancing at 100 % DR. We then loudness-balanced all stimuli at 80% and then 60% of the dynamic range. To do so each procedure started with the 90-pps stimulus at a level that was 80% or 60% of the way between its adaptive threshold and MCL in dB. Finally, the levels used for the other pulse rates presented in the temporal tasks were interpolated in dB using a linear regression. Note that, for rates other than 90 pps, the levels described here as corresponding to 60% and 80% DR are more accurately described as having the same loudness as a 90-pps pulse train at 60% or 80% DR. The shape of the loudness growth function may differ across pulse rates, and so the obtained values may not correspond exactly to 60% or 80% DR for each rate.

Finally we measured the low-rate RDR and upper limit using the same method as in experiments 1 and 2. This consisted of two adaptive runs of the RDR, followed by an MPC with 10 sub-blocks. This whole set of measures was obtained at loudness levels of 100%, 80%, and 60% of the DR in that order, and then repeated in reverse order. There was no effect of order, either within a set of measures at one loudness levels or across the two repeats of the loudness levels. We therefore report values obtained from the mean of all runs – a total of four adaptive RDRs and two sets of MPCs each obtained from 10 sub-blocks.

### B. Results

The level differences, relative to 100% DR, used for each subject are shown in Fig. 11 for the 80% - and 60%-DR conditions. Averaged across rates and subjects, these values were 1.17 and 2.34 dB for 80% and 60 % DR, respectively. A two-way RMANOVA (%DR X pulse rate) revealed a highly significant effect of %DR (F(1,11)=214.8 p<0.0005) and a significant effect of pulse rate (F(3,33)=4.06, p = 0.048). Inspection of Fig. 11 reveals that this latter effect was due to the level difference between 100% and 60% DR being slightly larger at 90 and 981 pps than at 162 and 399 pps. This was not true for the level difference between 100% and 60%, and so there was a significant interaction (F(3,33)=4.55, p<0.02). One subject, ME5, was tested on two electrodes and his data were treated as coming from separate subjects for the purposes of the RMANOVA.

**11).**
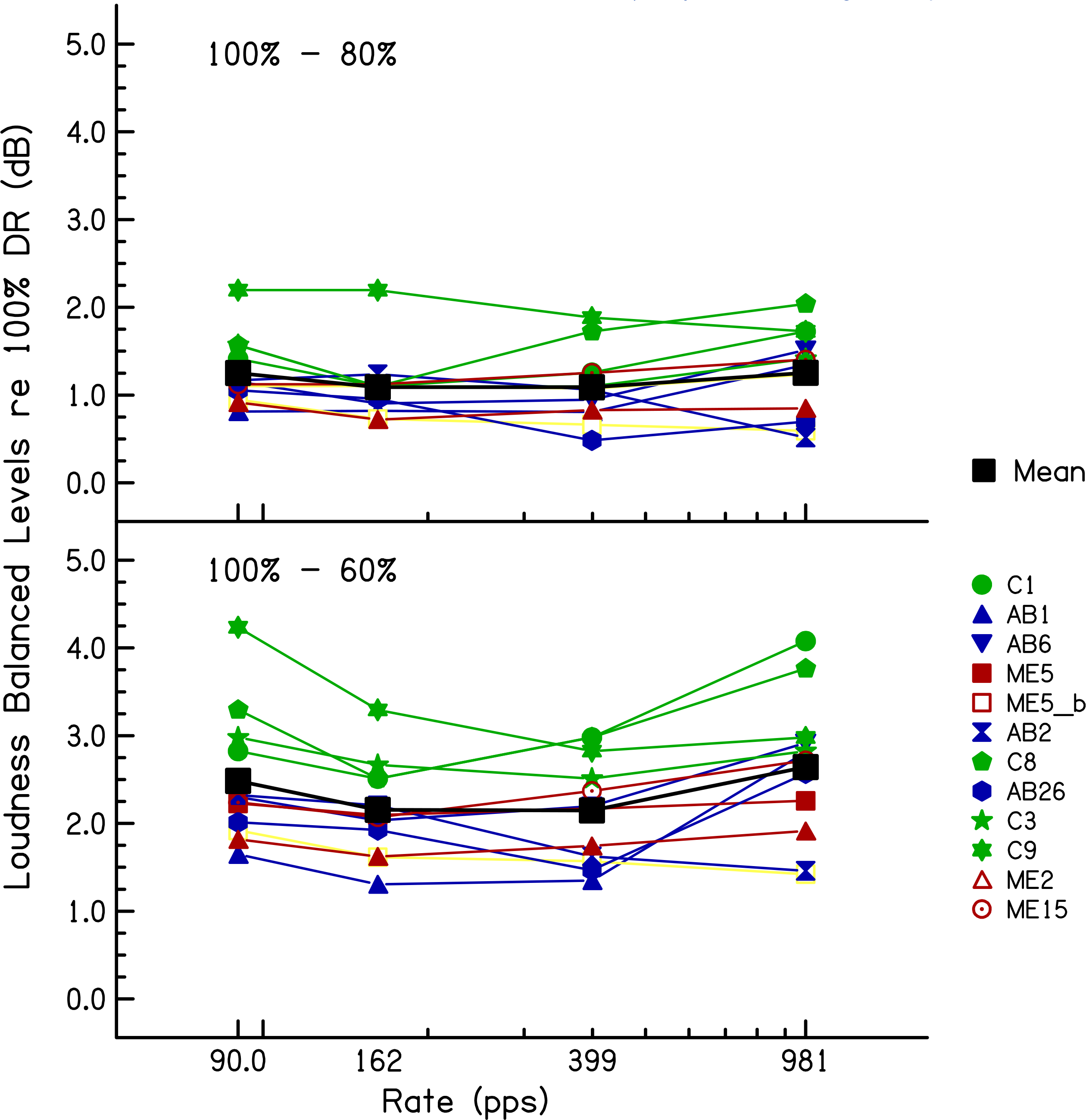
Loudness-balanced levels used in experiment 3, relative to 100% DR, for the 80%-DR (top) and 60%-DR (bottom) conditions. Levels are plotted as a function of pulse rate. Colored lines and symbol show data for individual listeners. The solid black squares connected by black lines show the mean levels across listeners.

Two aspects of the level differences are relevant to the interpretation of experiments 1 and 2. First, the average differences of 1.17 and 2.34 dB at 80% and 60% DR, relative to 100% DR, straddle the level difference of 1.7 dB between the first two sessions of experiment 1. Second, a comparison of the levels obtained from the more rigorous loudness balancing method used here, compared to that obtained by simply measuring MCLs, provides an estimate of the validity of the latter method, which was the only one used in experiments 1 and 2. We performed a two-way repeated-measures ANOVA (method X rate) on the levels obtained in experiment 3, excluding the data at 90 pps which, due to the procedure used, were identical for the two methods. This revealed a highly significant effect of rate, which unsurprisingly reflects the lower levels needed at higher rates (F(2,22)=30.7, p<0.001). The effect of method, however, just failed to reach significance (F(1,11)=4.6, p=0.054). Importantly, there was no interaction between method and rate, which would have meant that the loudness matching across rates depended on the method (F(2,22)=1.2, p=0.31)). Hence there was no evidence for a systematic difference in the results obtained with the two methods. As a test of reliability, we measured the average absolute difference between the two methods, again excluding the data at 90 pps. The mean absolute differences between the two methods were, at 162, 399, and 981 pps, 0.34, 0.35, and 0.56 dB, respectively. Furthermore, when averaged across rates, the MCLs and balanced levels correlated strongly across subjects (r=0.98, df=10, t=17.8, p<0.0001).

The low-rate RDRs are plotted as a function of level in Fig. 12 for all twelve data sets (11 subjects including subject ME5 who was tested on two electrodes). The mean data are shown by the thick black line. The effect of level differed across subjects but, on average, the RDR dropped from 1.17 to 1.13 to 1.11 as the level increased from 60 to 80 to 100% DR. This difference was statistically significant, as assessed by a one-way RMANOVA (F(2,22)=4.58, p=0.042); there was also a significant linear trend (F(1,11)=5.09, p=0.045). The increase in RDR from 100% to 60% DR corresponded to a factor of 1.04, with 95% confidence limits of 1.00-1.09.

**12).**
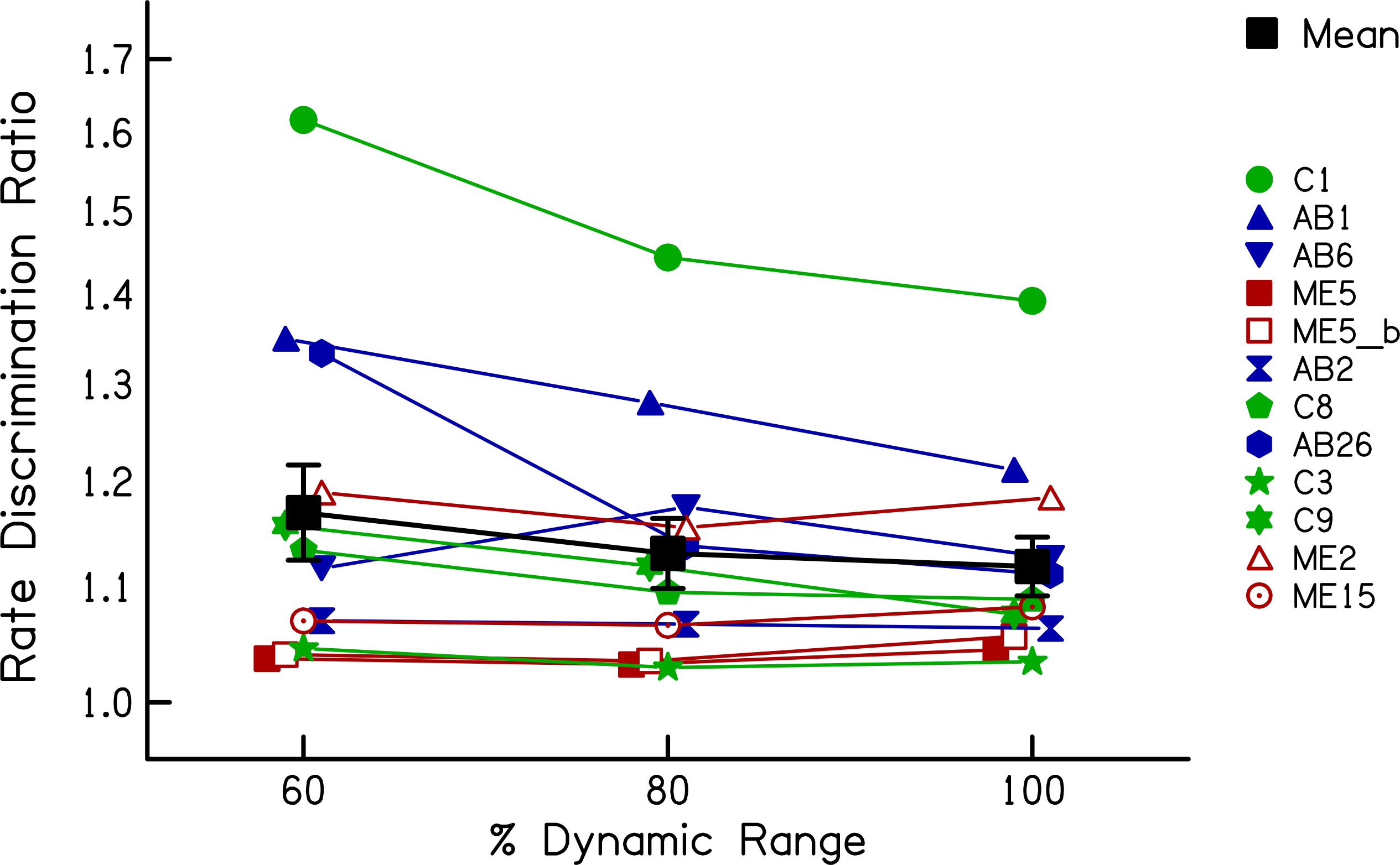
RDRs as a function of stimulus level expressed as a percentage of the dynamicrange, obtained in experiment 3. Coloured symbols show data for individuallisteners. The solid black line shows the geometric mean of all subjects.

The upper-limit data are shown in Fig. 13 using a similar format to Fig. 12. Here there is a much larger variation in the effect of level, which was not significant overall (F(2,22)=1.48, p=0.251). However it can be seen that one subject, ME5 (filled red squares), shows an unusually large *decrease* in the upper limit with increasing level. That subject was one of the three MedEl users tested on electrode 2, which is much more apical than any of the electrodes in the other devices. It was for this reason that the subject was re-tested on electrode 6. Those data, shown by the open red square, reveal a modest increase in the upper limit with increasing level, similar to that shown by many other subjects. Furthermore, one other MedEL user tested on electrode 2, subject ME2 (red triangles), showed a more modest decrease with increasing level. The other MedEl subject tested on electrode 2, ME15, showed no effect of level on the upper limit. Post-operative X-rays revealed insertion into the second turn of the cochlea for all three MedEL patients, with no evidence of any abnormal insertions in any of them. Possible reasons for these results are discussed in section 4.E. Whatever the explanation, it could be argued that the very apical location of MedEl electrode 2 represents a special case. In particular, for the purposes of using the level effects observed here to help interpret the results of experiments 1 and 2, we considered it worth re-analysing the upper limit data whilst excluding the three data sets obtained on MedEl electrode 2. This resulted in a highly significant effect of level (F(2,16)=15.33,p<0.001) and a significant linear trend (F(1,8)=20.09, p=0.02). It is illustrated by the dashed solid line, which shows the mean with those three data sets excluded. For this subset of data the mean upper limits were 315, 353, and 456 pps at 60, 80, and 100% RDR respectively. Hence the upper limits differed between 100% and 60% DR by a factor of 1.44 (95% confidence intervals: 1.120-1.75) and between 100% and 80% DR by a factor of 1.29 (1.14-1.47).

**13).**
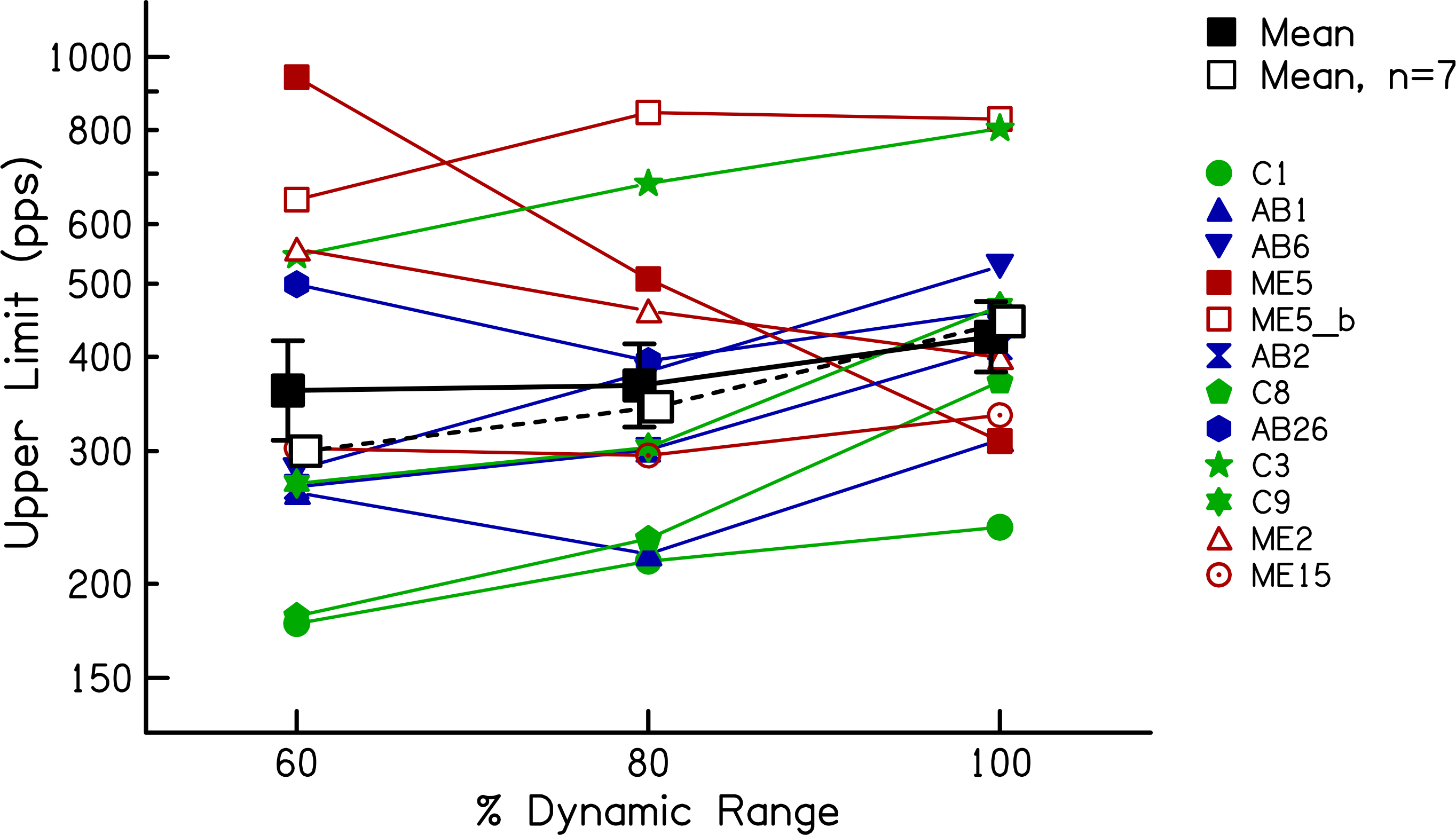
Upper limits as a function of stimulus level expressed as a percentage of the dynamic range, obtained in experiment 3. Coloured symbols show data for individual listeners. The solid black line shows the geometric mean of all subjects.

We also compared the effect on RDRs and the upper limit of the level differences used in this experiment with the differences observed between the first two sessions of experiment 1, where the MCL increased by 1.7 dB between sessions. As noted above, the 95% confidence intervals for the decrease in RDRs as the level was increased by (on average) 2.25 dB from 60% to 100% DR spanned factors of 1.00-1.23. This overlaps with the 95% confidence intervals for the change in RDR from session 1 to session 2, which spanned factors of 1.00-1.11. Similarly, the confidence limits for the increase in the upper limit from 60% to 100% RDR spanned factors of 1.20-1.75, which overlapped with the range of 1.20-1.53 observed in experiment 1 for the change between 0 and 2 months post activation.

## 4. DISCUSSION

### A. Does chronic stimulation affect temporal processing independently of level?

Experiment 1 showed a moderate but highly significant increase in the upper limit during the first two months post activation, and a significant reduction in the low-rate RDR over the same period. Because the measurements were obtained at MCL, which increased over the same period, it is important to consider whether the improvements in temporal processing were mediated by the change in physical stimulus level.

The additional analyses described in section 1.C provided some evidence that not all of the improvements in temporal processing were mediated by level differences. Here we re-visit that evidence in the light of the results of experiment 3. First, there was no between-subject correlation between the level-and upper-limit differences in sessions 1 and 2. However, experiment 3 showed that the effect of level on the upper limit could differ substantially across subjects. Hence, even if the upper-limit changes in experiment 1 were mediated by level, there may not have been a strong between-subject correlation. Second, the between-session level difference in experiment 1 was similar at all pulse rates, whereas the effect size was larger for the upper limit (which was presumably dominated by high rates) than for the low-rate RDR. This meant that, if level differences were responsible for the improvements between sessions 1 and 2, then a level change should have produced a larger effect size for the upper limit than for the low-rate RDR in experiment 3. This was not the case when the data from all subjects were included in the analysis (t(11)=0.18, p=0.57). If one excludes the data from the subjects tested on electrode 2 of the MedEl device, which yielded anomalous results for subjects ME5 and perhaps ME2, then the effect size is indeed numerically larger for the upper limit, but the difference just fails to reach significance (t(8)=1.70, p=0.06). Third, for a given subject, the change in MCL over the first three sessions did not always parallel the change in the upper limit. For example, the MCLs for subjects C105 and C106 increased between 2 and 6 months (Fig. 5) but their upper limits did not increase over this period (Fig. 3). However, experiment 3 showed that the effects of level on the upper limit are not always linear over the entire dynamic range; for individual subjects the upper limit could increase more from 60% to 80% DR than from 80% to 100% DR (e.g. subject C1), whereas for others the largest upper-limit change could be from 80% to 100% DR (e.g. subject C8). Overall, then, although experiment 1 showed that changes in MCL and in the upper limit did not correspond closely for all subjects, this cannot be taken as firm evidence against the hypothesis that the increase in upper limit was mediated by level changes. This was because level effects on the upper limit differ across subjects and across different portions of the dynamic range (DR). A similar conclusion applied to the changes in low-rate RDR, where the effect of level in experiment 3 also varied across subjects and across the DR.

Arguably stronger evidence that temporal processing improves, independently of level, arises from experiment 2. For the low-rate RDR, five subjects performed better when re-tested at 9 months than on the day of activation, despite the similar physical level and softer loudness in the latter session. A *caveat* is that, for this measure, performance improved during the first two sessions, and so one cannot exclude a procedural learning effect. This explanation is less likely for the one subject whose upper limit improved when re-tested at the same level after 9 months, because experiment 1 revealed no within-session effect for this measure. However this improvement was observed only for one subject, and so evidence that temporal processing improvements occur independently of level must be regarded as tentative. The possible influence of non-procedural learning effects are discussed further in section 4.D.

### B. Equating performance at equal level vs. equal loudness

Experiments 1 and 2 showed a significant and consistent increase in the upper limit when measured at approximately equal loudness (sessions 1 vs 2), but a less consistent improvement when measured at approximately equal level (sessions 1 vs 4). As noted above, this may mean that the observed improvements in temporal processing arise from, or are mediated by the same processes as, those responsible for the change in MCL over time. The interpretation of our findings, both from a practical and theoretical perspective, depends on the stage(s) of auditory processing at which those MCL changes occur. At one extreme, subjects may be feeling apprehensive on the day of activation and give loudness judgements that are greater than the “true” loudness. If that is the case then it would be appropriate to compare performance at equal physical levels, and we would conclude that there is no consistent improvement in temporal processing in the months following activation. At the other extreme, there could be changes at the level of the auditory nerve and/or its synapses with the cochlear nucleus, which effectively attenuate the input to higher stages of the auditory system. In that case, it would be more appropriate to equate performance at equal loudness, thereby roughly equating the input to higher auditory processes. Our results would then indicate that, for the same input, those processes had indeed become better at encoding temporal information. There are, of course a number of intermediate stages of processing at which the change in MCL may occur. From a theoretical perspective, a crucial distinction is whether the MCL changes occur at an earlier or later the stage of processing than that which limits performance on our temporal tasks.

Plasticity in the auditory system following implantation has been observed at very early stages of the auditory system. O’Neill *et al* (2010) performed histological analyses of the end bulbs of Held in normal-hearing (NH) cats and in congenitally deaf white cats that had either been implanted at 3 or 6 months, or were not implanted. The implanted cats were stimulated for 5 days a week for several months using a commercial-type processing strategy after which time they were sacrificed. The histology was roughly similar between the 3-month-implanted and NH cats on the one hand and the 6-month-implanted and unstimulated cats on the other, with the latter group having larger post-synaptic densities (PSDs) and a greater number of post-synaptic vesicles compared to the former. One unstimulated cat that was sacrificed after 3 months of stimulation showed similar results to those sacrificed after 6 months, suggesting that the histological changes were already complete after 3 months of stimulation, and that stimulation starting at 3 months restored the PSDs to normal, rather than preventing changes that would otherwise have occurred. Note that the chronic effect of implantation would likely have been to reduce the neural response to stimulation, and so may well have increased the MCL (in living animals). However it should be noted that these changes were observed in cats that had been deaf since birth, rather than in the adult-deafened human population observed here.

In tests with paediatric human CI patients, Gordon *et al* (2006) performed longitudinal measurements of the electrically evoked compound action potential (“ECAP”) and the electrically evoked auditory brainstem response (EABR) during the year following implantation. They observed reductions in the latency of the ECAP and of the EABR waves II, III and V during the first year of implant use, and also reported significant increases in inter-wave latencies. These measures were all obtained at equal loudness, but they additionally showed that the latency changes in the ECAP and in waves III and V remained significant when measured at equal level. Hence their results demonstrated changes at multiple stages of the auditory system, from the auditory nerve (ECAP) up to the inferior colliculus (change in wave V-wave II latency). However it is not known whether the latency reductions post implantation reflect a change in the accuracy of temporal encoding. Certainly, the size of the changes in ECAP latency, which were less than 50 μs, seems very small compared to the timing differences that our subjects detected in the present study. For example, the average low-rate RDR in session 2 (excluding C105 who performed very poorly) was 1.15, corresponding to a difference in inter-pulse interval of 1.22 ms. Similarly, the difference in inter-pulse-interval between two adjacent rates – 399 and 538 pps – tested in the pitch-ranking experiment was 647 μs.

Finally, it is worth noting that, from a practical perspective, it is not absolutely necessary to know the exact processing stage at which MCL changes occur, provided that the change in loudness is a genuine sensory effect rather than a change in response criterion. Suppose, for example, that the effective limit on temporal processing occurs at the IC, and that MCL changes are mediated by the auditory cortex. In theoretical terms, the response of the IC to any given stimulus has not changed in any way. However, from a practical perspective the more centrally mediated MCL changes would allow the listener to tolerate higher current levels, and this could in turn lead to the (higher level) stimulus being encoded with greater accuracy at the IC.

### C. “Temporally challenging” chronic stimulation

The stimuli that the subjects in experiment 1 heard following activation of their CI were the environmental and speech sounds of everyday life, processed via the ACE signal processing strategy. In common with most clinical processing strategies, such as Continuous Interleaved Sampling (“CIS”), ACE primarily conveys envelope information and discards temporal fine structure, and presents a constant rate of stimulation on each electrode. It is possible that the nature of the chronic stimulation and/or the need to make fine temporal judgements is important for any plasticity effects. A number of different types of chronic stimulation have been used, including 50-pps pulse trains (Argence et al., 2008; GABA inhibition in IC), a CIS-type processor (O’Neil et al., 2010; morphology of end bulbs of Held), 100-600-pps pulse trains (Middlebrooks, 2013; IC recordings), and 300-pps pulse trains amplitude-modulated at 30 Hz (Vollmer et al., 2005; Vollmer and Beitel, 2011; Vollmer et al., 2017; IC and auditory cortex recordings). These different forms of stimulation were tested with quite different measures, and so there is no firm evidence that one type of stimulation is more effective than any other. All we can say is that some plastic changes have been observed, for some measures, using forms of chronic stimulation that are no more “temporally challenging” than the ACE strategy in our subjects’ clinical processors.

### D. Learning and “delayed gains”

So far we have largely considered the between-session changes in the upper limit and in low-rate discrimination in terms of the effects of chronic stimulation and/or differences in level. However, performance on most behavioural tasks improves with practice, and a large body of literature has been devoted to the effects of training on perceptual tasks in both the visual and auditory domains. As a partial control for such effects we repeated both sets of measures within each session, on the assumption that at least some types of learning effect would be larger within than between sessions. This might include, for example, general familiarization with the testing set up and/or understanding what is meant by the term “pitch”. The finding that the upper limit increased across but not between sessions is, we think, more consistent with the between-session changes being due to the effects of chronic stimulation and/or level differences than to this type of learning. However, this may not apply to other types of learning. The classification of learning types differs across authors and studies, but typically distinguish between learning that is associated with the procedure, the task, or the stimulus used in training (e.g. Ortiz and Wright, 2009). These types are usually experimentally differentiated by measuring the generalization of training with a single procedure, task, and stimulus to other conditions that differ on one or more of those dimensions.

Of particular interest to the present study is that learning effects can appear many hours after training is complete. This has been observed, for example, for tasks such as the discrimination of inter-aural time differences (ITDs), consonant-vowel (CV) syllables, temporal intervals, and of frequency differences (Demany and Semal, 2002; Grimault et al., 2002; Ari-Even Roth et al., 2005; Ortiz and Wright, 2009, 2010). Usually the experiments studying these “delayed gains” are different from those that differentiate between the different types of learning, so it is not entirely clear which type or types of learning emerge over time. However, one study (Ortiz and Wright, 2010) trained different groups of subjects on either an ITD task with a 500-Hz sinusoidal signal, on an inter-aural level discrimination (ILD) task with a 4-kHz signal, or with no training. Both of the trained groups were then tested on an ITD task immediately after training and were better than the untrained group, and improved further when tested 10 hours later. This would appear to indicate a delayed gain for procedural learning, because it occurred both for a trained and an untrained task and stimulus. However, when tested after 24 hours the ITD-trained group improved further, but the ILD - trained group got worse. The authors suggested that stimulus-specific learning showed delayed gains up to 24 hours; the deterioration for the ILD-trained group is consistent with the effects of procedural learning wearing off.

Regardless of the type of learning involved, an important question for the interpretation of our results is whether learning of the pitch-ranking task was likely to have increased between sessions, even though no gains were obtained within sessions. Although we cannot rule this out, there are some differences between the paradigm used here and those usually used to study delayed gains that make this less likely. Perhaps the most important of these is that delayed gains are usually studied with extensive training on a single task in which feedback is given. In contrast, our pitch-ranking procedure involved no feedback and was mixed with the low-rate discrimination task. The low-rate discrimination task did involve feedback, and so one might expect greater learning on that task than on the pitch-ranking task. There may of course have been some transfer between tasks, but this could not explain why the between-session effect size was larger for the (no-feedback) upper-limit measure than for low-rate discrimination. Second, we recently used a very similar procedure to study the effects of a pharmaceutical agent on temporal discriminations by CI listeners (Carlyon et al., 2018). That study involved testing 12 CI users before and after two 28-day periods of taking either a drug or a placebo. No effect of the drug was observed. More importantly for the present discussion, the upper limit did not differ between the first and third sessions, which were the pre-drug and pre-baseline estimates, and so there was no indication of any between-session learning effects for that task, although some improvement was observed for the RDR. Finally, it is known that the time course of learning effects can differ substantially between different auditory tasks (Wright and Sabin, 2007). Most pitch-related learning studies have used pure-tone stimuli, which are quite different from the electric pulse trains presented to our subjects. However, one study measured learning effects for the discrimination of bandpass filtered harmonic complexes, with F0s such that the harmonics were resolved or unresolved by the peripheral auditory system (Grimault et al., 2000). This is of interest because of evidence that discrimination and pitch judgements of unresolved complexes provide a close parallel with that of electric pulse trains presented to CI listeners (McKay and Carlyon, 1999; Carlyon et al., 2002; van Wieringen et al., 2003). Grimault *et al* measured F0 discrimination for two sessions without feedback, followed by six training sessions with feedback. For the resolved group thresholds dropped as soon as feedback was introduced, and then showed a further gradual drop over the daily training sessions, as has been observed with pure-tone frequency discrimination. In contrast, thresholds for the unresolved group also dropped as soon as feedback was introduced, but no further drop occurred during training. For these stimuli, which are arguably closest to those used in the present experiment with CI listeners, the results are consistent with feedback informing subjects of the task requirements, but with no further learning or training effects occurring.

### E. Anomalous effects of level on the upper limit of temporal pitch

For most subjects in experiment 3, the upper limit increased with increasing stimulus level. However, subjects ME5 and possibly ME2 showed the opposite effect. These were two of the three subjects stimulated on an apical electrode of the MedEl device, which has a much longer array than those in the Advanced Bionics and Cochlear CIs. One possible explanation comes from the observation that so-called place-pitch reversals – whereby stimulating a more basal electrode results in a lower pitch – are more common in the apical third of the MedEL array than in the central or basal thirds (Kenway et al., 2015). The reversals might reflect either cross-turn stimulation or the particular cochlear/nerve anatomy at the apex, where Rosenthal’s canal has ended and where nerve fibers are tightly packed. It may be that increasing stimulus level also increases the proportion of responding neurons that have CFs remote from those closest to the stimulating electrode, and that this somehow introduces place-pitch confounds that reduce the upper limit. To do so they would either have to reduce the place pitch with increasing rate, or perhaps simply confuse the subject so that s/he responds in a less consistent fashion throughout each block of trials.

### F. Summary and implications

We measured thresholds for the discrimination of low-rate pulse trains (RDRs) and the upper limit of pitch on the day the implant was activated and at 2 and 6 months later. Performance on both tasks showed a high across-subject correlation between the first two sessions. The two tasks did not correlate with each other, consistent with the results of Cosentino *et al* (2016) and with the conclusion that the marked differences observed across subjects in each task are not dominated by the same limitation. Performance improved, on average, between the first two sessions, with the RDR decreasing from 1.23 to 1.16 (after one poor-performing subject’s data were excluded) and the upper limit increasing from 357 to 485 pps. There were no significant changes between 2 and 6 months. These results place an upper bound on the amount of improvement in temporal processing that occurs post implantation in adult-deafened human CI listeners, and show that any improvements occur within the first two months. They provide no evidence for the idea that the upper limit decreases as a result of chronic exposure to a signal-processing strategy that discards TFS information.

The MCLs, and hence the testing level, increased on average between 0 and 2 months post-activation, and for some subjects increased further between 2 and 6 months. At least some of the improvement in temporal processing may have been due to this level increase, because performance on both tasks generally improved with increasing level when tested in a separate group of subjects. No further evidence for this explanation was found; for example the changes in MCL did not correlate with changes in the upper limit or RDRs, and some subjects whose MCLs increased between 2 and 6 months did not show a change in the upper limit or RDR. When a subset of subjects were re-tested at 9 months, using similar stimulus levels to those observed at 0 months, the pattern of changes in performance were mixed. Some subjects did show significantly improved performance at 9 months, despite the stimuli sounding softer. Others showed significantly worse performance, indicating that reduced loudness can impair performance even when the stimulus level is the same.

Overall it proved difficult to disentangle the effects of differences in MCL and loudness on temporal processing. It is worth noting that a related complication may apply to the physiological data on temporal processing obtained from single-unit recordings in animals. This is because those recordings are usually obtained at a stimulus level that is, for example, a fixed amount above each neuron’s threshold. It is possible that deprivation affects those thresholds, which in turn affects the stimulus level needed to record the responses. Such effects would likely not be noticed, because the single-unit recordings necessarily involve comparisons between animals, rather than a longitudinal comparison such as that used here, and because overall sensitivity differs markedly across animals (as it does across humans). In both cases, one could argue that plasticity has affected temporal processing, either directly or indirectly via the changes in overall sensitivity.

Regardless of the link between the level and temporal-processing changes observed here, the present article provides new information on the plasticity of the adult auditory system. In addition it raises the possibility that the improvements in speech perception that are observed in the months following implantation may not be entirely due to learning, but may instead reflect changes in sensory processes that can be observed using psychophysical techniques and with simple stimuli. A possible practical application is that knowledge of these sensory changes may help explain the substantial across-listener differences in the improvement in speech perception following implantation.

